# The emergence of a novel CA1 spatial map requires direct entorhinal input

**DOI:** 10.64898/2026.07.02.736055

**Authors:** Oscar M. T. Chadney, Matteo Guardamagna, Felix Dörr, Lucie A. L. Descamps, Federico Stella, Francesco Battaglia, Clifford G. Kentros

**Affiliations:** Kavli Institute for Systems Neuroscience and Centre for Algorithms in the Cortex, Norwegian University of Science and Technology, Trondheim, Norway; Donders Institute for Brain, Cognition and Behaviour, Radboud University, Nijmegen, Netherlands; Institute of Neuroscience, University of Oregon, Eugene OR, USA

**Keywords:** Hippocampus, CA1, entorhinal cortex, place cell, spatial coding, novelty, remapping

## Abstract

The hippocampus and its main input, the entorhinal cortex, are essential for memory formation, yet how they interact to do so remains unresolved. Hippocampal CA1 pyramidal neurons exhibit spatial receptive fields which reorganize unpredictably upon exposure to a novel environment, a process called remapping, considered a model of memory formation. CA1 neurons integrate both hippocampal and direct entorhinal inputs, but their relative contributions to spatial coding are unclear. Here we combined population recordings of CA1 neurons with optogenetic silencing of their direct entorhinal input. While this manipulation did not impact spatial firing in a familiar environment, it impaired remapping in a novel environment, resulting in the emergence of a stable hybrid map combining features of both environments. This shows the direct entorhinal pathway plays a specific role in CA1 novelty detection, enabling the plasticity necessary to drive the reorganisation of spatial firing patterns, preventing interference between memories in different contexts.

## Introduction

The ability to register novel information in our surroundings is fundamental to forming memories. The hippocampal formation is central to this process, supporting the encoding and retrieval of episodic experiences ^1–6^. Hippocampus place cells exhibit spatially selective firing ^7^ which at the population level forms a representation of self-location that underlies a cognitive map ^2,8,9^. When an animal encounters a novel environment, place cell firing patterns unpredictably reorganise in a process known as *remapping* ^10–12^, forming orthogonal environmental representations that support accurate context-specific spatial memory ^13,14^. However, the circuit mechanisms driving remapping remain unclear.

CA1 pyramidal neurons, the principal output of the hippocampus, receive two major inputs originating from the entorhinal cortex (EC): indirectly via the tri-synaptic pathway (EC layer II – dentate gyrus – CA3 – CA1) targeting proximal dendrites, and the direct pathway (EC layer III (ECIII) – CA1) ^15,16^ projecting to distal dendrites. This dual input has attracted considerable interest regarding the roles of both inputs in memory encoding and retrieval ^17–24^. Experimental evidence and numerous models suggest that remapping arises from the interplay between a dentate gyrus novelty signal ^25^, grid-to-place transformation ^21,26–34^ and attractor dynamics in CA3^35^. However, several observations challenge a model in which CA1 remapping is solely inherited from the tri-synaptic pathway: CA1 can form stable representations prior to CA3^13,36^and can reorganise independently during learning ^37^, suggesting that the emergence of novel representations in CA1 engages local network computations ^38^.

The direct pathway can drive dendritic plateau potentials, that enable nonlinear integration of CA3 and ECIII inputs, implicated in place field formation ^39,40^. Despite this mechanistic insight, the role of the direct entorhinal input in CA1 population-level remapping during the exploration of a novel environment remains unknown.

Here, we combined population recordings of CA1 place cells with targeted optogenetic silencing of direct ECIII inputs during exploration of familiar and novel environments. This approach preserved ECIII somatic activity and the tri-synaptic pathway, allowing us to study the specific contribution of the direct entorhinal-hippocampal pathway in shaping novel CA1 spatial maps. Our results provide direct circuit-level insight into how the hippocampus encodes and distributes new knowledge about the surrounding environment.

## Results

### Direct entorhinal input inhibition does not affect a familiar CA1 spatial map

To transiently inhibit direct entorhinal input to CA1, we used a transgenic mouse cross ^41^ expressing the inhibitory opsin JAWS ^42^ specifically in ECIII cells ^43^ (Figure 1A). An optic fiber implanted above CA1 targeted ECIII axon terminals (Figure 1B and Figure S1A), enabled local inhibition of the direct entorhinal inputs. Animals expressing the inhibitory opsin (Opsin group) were compared to littermate controls (Control group) that received the same implants, light-stimulation and behaviour protocols (Figure S1A-E and Methods). CA1 place cells were recorded while animals were exposed to a familiar environment, first without (F1) then with inhibition of the direct entorhinal input (F2, Figure 1C).

**Figure 1.**
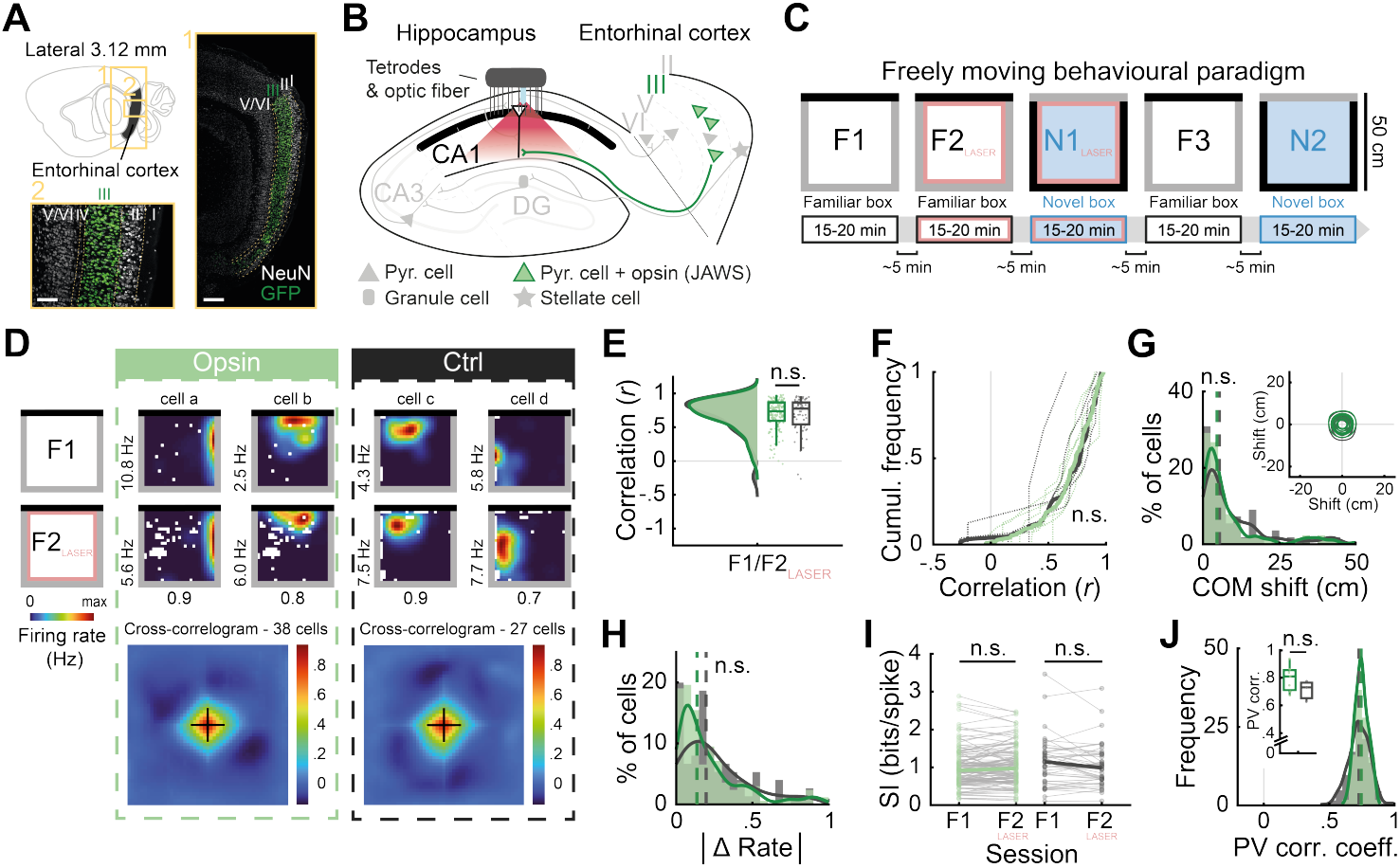
Direct entorhinal input inhibition does not affect a familiar CA1 spatial map. (A)Fluorescent *in situ* hybridization of a sagittal section showing the JAWS-GFP transgene (green) confined to ECIII neurons. NeuN, white. Scale bars, left: 150µm; right: 250µm. **(B)** Schematic illustrating CA1 population recordings and optogenetic inhibition of ECIII projections. **(C)** Behavioural paradigm. Animals explored a familiar environment without (F1) and with ECIII input inhibition (F2), followed by the exposure to a novel environment with inhibition (N1) before re-exposure to the familiar (F3) and novel (N2) environments with intact entorhinal input. **(D)** Top: rate maps from two CA1 place cells per group during F1 and F2. Peak firing rates are bottom left; correlation coefficients between F1 and F2 are shown below. Bottom: cross-correlograms of simultaneously recorded place cells from representative animals. **(E)** Pearson’s correlations between F1 and F2 place cell rate maps. Dots represent individual cells (Opsin, *N*=156; Control, *N*=65). *p*=0.7248, Z=-0.3521, Wilcoxon rank-sum test. **(F)** Cumulative frequency diagram of single-cell rate map correlations (Opsin, *N*=156; Control, *N*=65). Dashed lines, individual animals (Opsin, *n*=7; Control, *n*=4); solid lines, pooled data. *p*=0.8106, D=0.0923, Kolmogorov-Smirnov test. **(G)** Distribution of place field center-of-mass (COM) shifts (Opsin, *N*=127; Control, *N*=55). *p*=0.3105, Z=1.0142, Wilcoxon rank-sum test. Inset: contour plot showing direction and distance of place field displacement for representative Opsin (*N*=32) and Control (*N*=25) animals. **(H)** Distribution of place cell absolute firing rate change (Opsin, *N*=168; Control, *N*=75). *p*=0.1188, Z=-1.5598, Wilcoxon rank-sum test. **(I)** Spatial information (SI) of place cells. Dots represent individual cells. Opsin (*N*=113): *p*=0.1557, Z=-1.4195. Control (*N*=44): *p*=0.0669, Z=1.8322. Wilcoxon signed-rank test. **(J)** Distribution of population vector (PV) correlations. Inset: animal-averaged correlations (Opsin, *n*=7; Control, *n*=3). *p*=0.2066, permutation test. Box plots show median and interquartile range. Dashed lines overlaying histograms show medians. Green, Opsin; gray, Control.

In line with previous findings ^43^, both Opsin and Control groups maintained stable spatial tuning across familiar environment sessions with and without light stimulation (Figure 1D-1J). Cross-correlograms of simultaneously recorded place cells displayed a clear peak at the origin (Figure 1D, bottom), indicating stable spatial maps across sessions. Indeed, all single-cell measures showed no significant difference between opsin and control animals (Figure 1E-1I). Likewise, the population vector correlation, a population-level means to assess the similarity of place cell activity, revealed no significant difference between groups (Figure 1J). These results indicate that silencing the direct entorhinal-CA1 projection does not affect the recall or maintenance of the spatial map of a familiar environment.

### Direct entorhinal input is required for the emergence of a novel CA1 spatial map

We next explored the role of the direct ECIII input in generating a novel CA1 spatial map by inhibiting ECIII axon terminals during exposure to a novel environment (N1, Figure 1C and Figure 2). Control animals reliably exhibited robust global reorganisation of spatial firing patterns in the novel environment (N1, Figure 2A-2G), evidenced by the lack of peak at the origin of the cross-correlogram (Figure 2A, bottom right). Strikingly, remapping was inhibited in Opsin animals (Figure 2A-2I), as can be seen in the residual peak at the origin of the cross-correlogram (Figure 2A, bottom left), suggesting persistent similarity to the familiar environment map. Similarly, Pearson’s correlations between familiar (F2) and novel (N1) environment rate maps were significantly higher in Opsin animals (Figure 2B and 2C) and place fields shifted significantly less (Figure 2D), while the percentage of place cells remained the same (Figure S2B). This effect could not be attributed to differences in spike sorting quality ^44,45^, nor behaviour (Figure S1D and S1E). A larger fraction of Opsin place cells retained high similarity across environments (23.4% with *r* >0.5) compared to Controls (8% with *r* >0.5) (Figure S2A), yielding a hybrid map combining familiar and novel representations. Changes in firing rate upon exposure to the novel environment were comparable between groups (Figure 2E), and for both groups, significantly higher than between consecutive familiar sessions (Figure S2C and S2D), indicating that Opsin place cells can undergo substantial firing rate changes despite minimal spatial reorganization ^46^. While spatial information of Control place cells was similar in the familiar and novel environments, it decreased in Opsin place cells (Figure 2F), likely reflecting increased place field size (Figure S2E) rather than out-of-field firing (Figure S2F). At the population level, vector correlations between familiar and novel environments were significantly higher in Opsin animals (Figure 2G). Together, these results demonstrate that direct entorhinal input to CA1 plays a key role in the emergence of an orthogonal spatial map of a novel environment.

**Figure 2.**
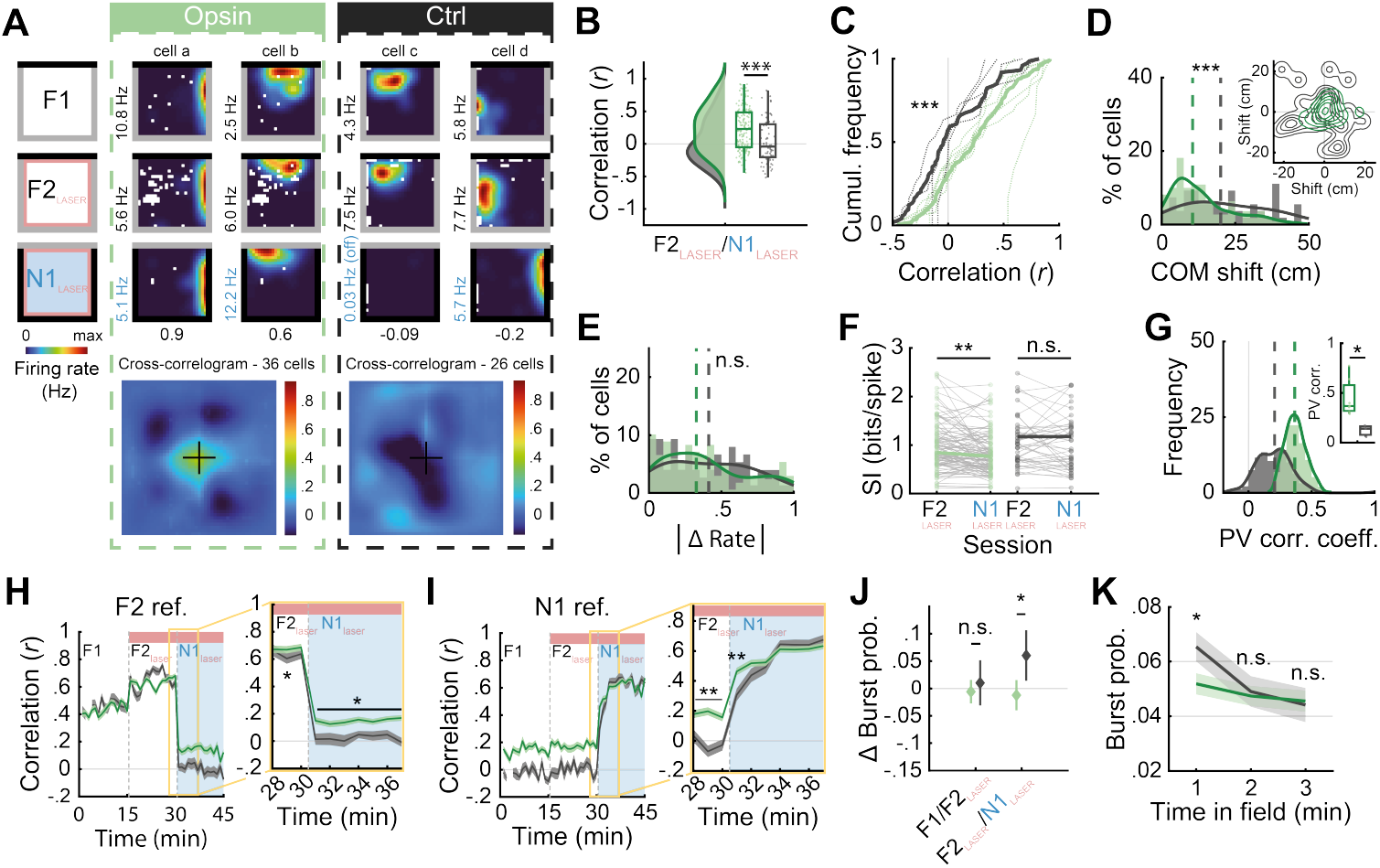
Direct entorhinal input is required for the emergence of a new CA1 spatial map. **(A)** Top: rate maps from two CA1 place cells per group during F1, F2 and N1. Peak firing rates are bottom left; correlation coefficients between F2 and N1 are shown below. Bottom: cross-correlograms of simultaneously recorded place cells from representative animals. **(B)** Pearson’s correlation between F2 and N1 rate maps. Dots represent individual cells (Opsin, *N*=165; Control, *N*=74). \*\*\**p*=3.6×10^−5^, Z=4.1341, Wilcoxon rank-sum test. **(C)** Cumulative frequency diagram of single-cell rate map Pearson correlation coefficients (Opsin, *N*=165; Control, *N*=74). Dashed lines, individual animals (Opsin, *n*=7; Control, *n*=4); solid lines, pooled data. \*\*\**p*=5.7×10^−4^, D=0.2775, Kolmogorov-Smirnov test. **(D)** Distribution of place field center-of-mass (COM) shifts (Opsin, *N*=110; Control, *N*=54). \*\*\**p*=1.7×10^−4^, Z=3.7598, Wilcoxon rank-sum test. Inset: contour plot showing direction and distance of place field displacement for representative Opsin (*N*=30) and Control (*N*=25) animals. **(E)** Distribution of place cell absolute firing rate change (Opsin, *N*=168; Control, *N*=75). *p*=0.6249, Z=-0.4890, Wilcoxon rank-sum test. **(F)** Spatial information (SI) of place cells. Dots represent individual cells. Opsin (*N*=99): \*\**p*=0.0013, Z=3.2076. Control (*N*=45): *p*=0.6075, Z=0.5136. Wilcoxon signed-rank test. **(G)** Distribution of population vector (PV) correlations. Inset: animal-average correlations (Opsin, *n*=7; Control, *n*=3). \**p*=0.0331, permutation test. **(H)** Left: Pearson’s correlation between 1-minute block rate maps (sessions F1, F2 and N1) and the entire familiar session rate map (F2 reference, ref.) (Opsin, *N*=168; Control, *N*=75). Right: expanded transition from the familiar (F2) to the novel (N1) environment. All panels show medians±SEM. See Table S1 for statistics. **(I)** Same as **(H)**, using the novel session (N1) as the reference. See Table S1 for statistics. **(J)** Change in burst probability of place cells (median±SEM) between consecutive familiar sessions with (F1) and without (F2) ECIII input inhibition (Opsin, *N*=158; Controls, *N*=67, *p*=0.2182, Z=0.7783, Wilcoxon rank sum test) and between consecutive familiar (F2) and novel (N1) sessions with ECIII inhibition (Opsin, *N*=161; Controls, *N*=69, \*\**p*=0.0083, Z=-2.3961, Wilcoxon rank-sum test). **(K)** Burst probability during the first three minutes within place field (N1) (Opsin, *N*=168; Control, *N*=75; mean±SEM). First minute: \**p*=0.035, Z=-2.1085; second: *p*=0.9356, Z=-0.808; third: *p*=0.6485, Z=0.4558, Wilcoxon rank-sum test. Box plots show median and interquartile range. Dashed lines overlaying histograms show medians. Green, Opsin; gray, Control.

To determine how blocking the direct ECIII input affects the kinetics of remapping we correlated one-minute rate maps versus the rate map from either the entire familiar (F2) or novel (N1) environment sessions to capture the decorrelation from the familiar map (Figure 2H) and the emergence of the novel map (Figure 2I), respectively. Correlation with the familiar map remained high throughout consecutive familiar environment sessions (F1, F2) in both groups (Figure 2H, left), but rapidly dropped within the first minute in the novel environment and was thereafter stable (Figure 2H and S3A). However, this decorrelation was complete in Controls, but only partial in Opsin animals (Figure 2H). In contrast, the novel map emerged progressively over several minutes in both groups, eventually reaching correlation levels resembling the familiar environment (Figure 2I). The minute-by-minute change in correlation, was significantly greater and laster longer in the Control group (Figure 2I), reflecting a larger proportion of remapping cells.

At a mechanistic level, conjunctive CA3 and ECIII input can generate dendritic plateau potentials which produces burst firing in CA1 pyramidal cells ^39,47^, processes thought to underlie place field formation. Consistent with this interpretation, during the first exposure to the novel environment (N1), burst probability increased in the Control group as expected ^48^, but remained unchanged in the Opsin group (Figure 2J). This difference was confined to the first minute within the place field location (Figure 2K), emphasizing a critical early time window for direct entorhinal input in the novel space. The effect was specific to the novel environment, as ECIII input inhibition did not affect burst probability in the familiar environment (Figure 2J). This absence of increased burst firing may limit synaptic plasticity, constraining CA1 representations to the familiar map.

### The emergence of a novel CA1 map relies on direct entorhinal input during the first instances of exploration

Having prevented global remapping during the first exposure to the novel environment (N1), we hypothesised that an orthogonal CA1 spatial map would emerge upon re-exposure with intact ECIII input (N2, Figure 1C). Surprisingly, CA1 spatial firing patterns of the Opsin group during the second exposure (N2) remained highly correlated to those of the first exposure under ECIII input inhibition (N1, Figure 3A-3G), indicating the persistence of the previously formed hybrid representation. Single-cell and population measures showed no significant differences between groups (Figure 3B-3G). Moreover, during re-exposure (N2) both decorrelation from the familiar map and reactivation of the hybrid map occurred within the first minute (Figure S3A and S3B), in contrast to the gradual emergence of a new map during initial exposure (N1, Figure 2I). Thus, the hybrid map formed during ECIII input inhibition behaves as a stable spatial map rather than undergoing remapping.

**Figure 3.**
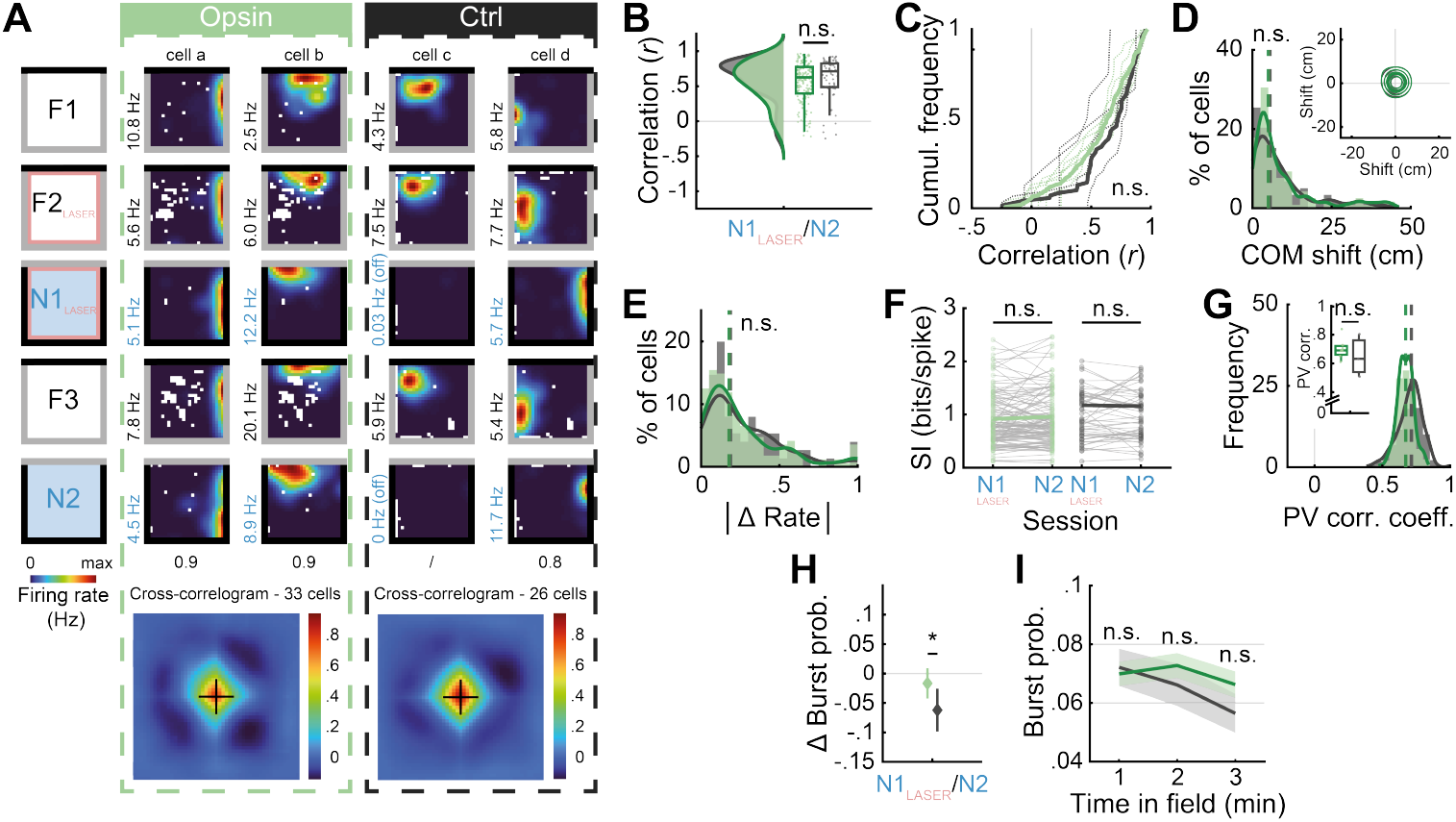
The hybrid CA1 spatial map persists despite intact direct entorhinal input. **(A)** Top: rate maps from two CA1 place cells per experimental group during all sessions. Peak firing rates are bottom left; correlation coefficients between N1 and N2 are shown below. Bottom: cross-correlograms of simultaneously recorded place cells from representative animals. **(B)** Pearson’s correlation between N1 and N2 rate maps. Dots represent individual cells (Opsin, *N*=155; Control, *N*=70). *p*=0.0878, Z=-1.7071, Wilcoxon rank-sum test. **(C)** Cumulative frequency diagram of single-cell rate map Pearson correlation coefficients (Opsin, *N*=155; Controls, *N*=70). Dashed lines, individual animals (Opsin, *n*=7; Control, *n*=4); solid lines, pooled data. *p*=0.0666, D=0.1846, Kolmogorov-Smirnov test. **(D)** Distribution of place field center-of-mass (COM) shifts (Opsin, *N*=110; Control, *N*=54). *p*=0.7616, Z=-0.3034, Wilcoxon rank sum test. Inset: contour plot showing direction and distance of place field displacement for representative Opsin (*N*=27) and Control (*N*=25) animals. **(E)** Distribution of place cell absolute firing rate change (Opsin, *N*=168; Controls, *N*=75). *p*=0.4775, Z=-0.7102, Wilcoxon rank-sum test. **(F)** Spatial information (SI) of place cells. Dots represent individual cells. Opsin (*N*=116): *p*=0.5536, Z=-0.5923. Control (*N*=54): *p*=0.6389, Z=0.4693. Wilcoxon signed-rank test. **(G)** Distribution of population vector (PV) correlations. Inset: animal-average correlations (Opsin, *n*=7; Controls, *n*=3). *p*=0.4380, permutation test. **(H)** Change in burst probability of place cells (median±SEM) between the first (N1) and second (N2) exposure to the novel environment with and without inhibition, respectively (Opsin, *N*=157; Control, N*=*68). \**p*=0.0352, Z=1.8098, one-sided Wilcoxon rank-sum test. **(I)** Burst probability during the first three minutes within place fields (N2) (Opsin, *N*=168; Controls, *N*=75; mean±SEM). First minute: *p*=0.7148; second: *p*=0.3211; third: *p*=0.2535, Wilcoxon rank-sum test. Box plots show median and interquartile range. Dashed lines overlaying histograms show medians. Green, Opsin; gray, Control.

Similarly, we predicted that reinstating ECIII input would restore increased burst firing in Opsin animals (N2), as observed in Control animals during the first exposure (N1). In Controls, burst probability decreased in the second exposure (N2) relative to the first (N1, Figure 3H), consistent with familiarization to the environment. In contrast, burst probability remained unchanged in Opsin animals (Figure 3H and 3I). Taken together, these results demonstrate that direct entorhinal input is critical during the initial moments in a novel environment, driving the formation of an orthogonal map. Once this time window has passed, subsequent reorganisation is strongly constrained, dissociating the generation of a place field from its consolidation.

### Independent CA3 dynamics during inhibition of direct entorhinal input

We next asked whether the lack of remapping observed in CA1 during ECIII terminal inhibition reflects a CA1-specific effect or arises from upstream changes in CA3. To address this, we performed simultaneous population recordings of CA1 and CA3 place cells in a subset of Opsin animals subjected to the same experimental paradigm (Figure 4A). In the familiar environment, like CA1, CA3 place cells maintained stable spatial coding despite ECIII input inhibition (Figure 4B and 4C, left), with rate map correlations (F1vsF2) significantly higher than CA1 place cells (Figure 4B, left), consistent with previous findings ^36^. Upon exposure to the novel environment (N1), in contrast to CA1 neurons, CA3 place cells underwent robust remapping (Figure 4B and 4C, middle), with rate map correlations between the familiar and novel environments (F2vsN1) significantly lower than CA1 place cells (Figure 4B, middle). Thus, while CA1 remapping was impaired under ECIII input inhibition, CA3 representations reorganised normally. During re-exposure to the novel environment without inhibition (N2), both CA1 and CA3 spatial maps were stable and highly correlated with those formed during the first exposure (N1) (Figure 4B, right panel), indicating their consolidation.

**Figure 4.**
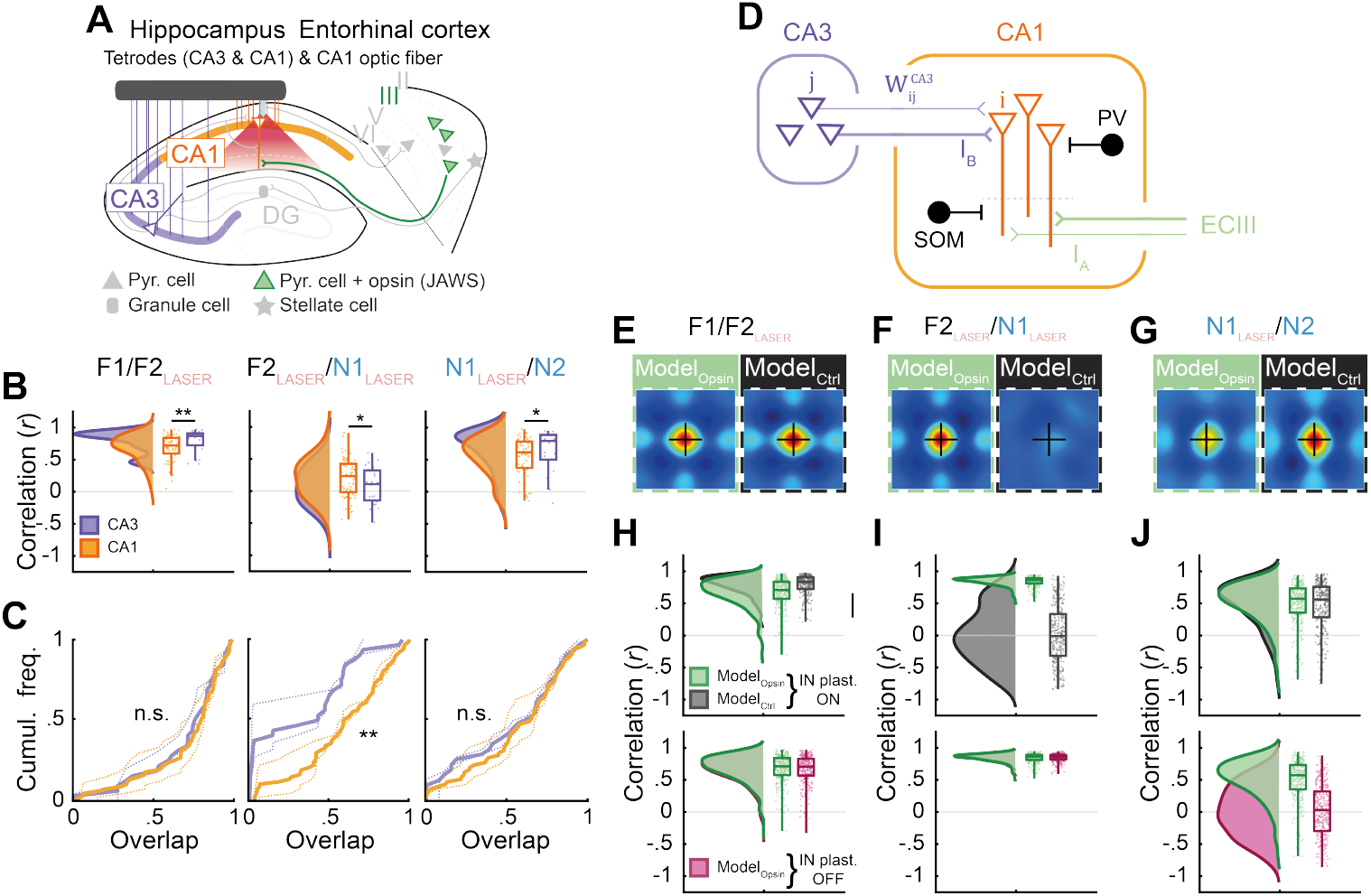
A dual-input model for CA1 spatial map formation. **(A)** Schematic illustrating optogenetic inhibition of ECIII projections in CA1 combined with simultaneous CA1 and CA3 population recordings. **(B)** Pearson’s correlations of place cell rate maps from simultaneous CA1 and CA3 recordings (*n*=2). Left: F1 versus F2 (Opsin CA1, *N*=60; Opsin CA3, *N*=23). \*\**p*=0.035, Z=-2.7013, one-sided Wilcoxon rank-sum test. Middle: F2 versus N1 (Opsin CA1, *N*=66; Opsin CA3, *N*=25). \**p*=0.0411, Z=1.7382, one-sided Wilcoxon rank-sum test. Right: N1 versus N2 (Opsin CA1, *N*=63; Opsin CA3, N=19) \**p*=0.0392, Wilcoxon rank-sum test. **(C)** Cumulative frequency diagrams of single-cell firing rate overlap (Opsin CA1, *N*=67; Opsin CA3, *N*=32). Dashed lines, individual animals (*n*=2; simultaneous CA1 and CA3 recordings); solid lines, pooled data. Left: F1 versus F2. *p*=0.9790, D=0.0984, Kolmogorov-Smirnov test. Middle: F2 versus N1. \*\**p*=0.0031, D=0.3750 Kolmogorov-Smirnov test. Right: N1 versus N2. *p*=0.7790, D=0.1387, Kolmogorov-Smirnov test. **(D)** Spiking neural-network model of the hippocampal circuit. CA1 pyramidal neurons are modeled as two-compartment Leaky Integrate-and-Fire neurons receiving basal CA3 input (IB) and apical ECIII input (IA). Single-compartment somatostatin (SOM) and parvalbumin (PV)-like interneurons (black) provided apical and basal compartment inhibition, respectively. **(E-G)** Cross-correlograms of simulated place cell activity for Opsin (Model_Opsin_, *N*=289 place cells) and Control (Model_Ctrl_, *N*=289 place cells) conditions between sessions F1 and F2 **(E)**, F2 and N1 **(F)**, and N1 and N2 **(G). (H-J)** Top: Pearson’s correlations of simulated place cell rate maps for Opsin and Control conditions with inhibitory plasticity enabled (IN plast. ON). Bottom: Pearson’s correlations of simulated place cell rate maps for Opsin conditions with (green) and without (pink) inhibitory plasticity enabled. Comparisons are shown for sessions F1 and F2 **(H)**, F2 and N1 **(I)**, and N1 and N2 **(J)**. Dots represent individual simulated cells (Model_Opsin_, *N*=289; Model_Ctrl_, *N*=289). Box plots show median and interquartile range. Orange, Opsin CA1; purple, Opsin CA3.

These findings support a model in which the emergence of a novel CA1 spatial map arises from local integration of CA3 and direct entorhinal inputs.

### A dual-input model for CA1 spatial map formation

To probe the circuit mechanisms underlying CA1 remapping, we built a spiking network model of the hippocampal circuit (Figure 4D). A population of two-compartment CA1 neurons received convergent CA3 input via feed-forward plastic weights (Schaffer collaterals, SC) targeting the basal compartment and direct ECIII input targeting the apical compartment. During two-dimensional virtual exploration, both pathways provided place-specific activity, and we hypothesized that CA1 bursting reflects mismatch between CA3 and ECIII inputs.

We reproduced the experimental paradigm *in silico*. The network was first trained in a familiar environment with intact ECIII input and SC plasticity enabled to establish CA3→CA1 associations. Novel exposure (N1) was simulated by shuffling CA3 and ECIII place-field centers while preserving a small but non-zero overlap with the familiar map (*r* ≈ 0.2), consistent with experimental observations that CA3 rate maps remain weakly but significantly correlated between familiar and novel environments (Figure 4B, middle, \*\*\**p*<0.001, permutation test) and with reports describing stable and dynamic coding in CA3 during novelty ^49^. Control and Opsin conditions were modeled by maintaining or silencing ECIII input, respectively, and remapping was quantified by comparing CA1 spatial activity across sessions (Figure 4E–4J).

CA1 remapping arose from three interacting learning rules: (i) burst-dependent SC plasticity, (ii) Hebbian-like SC plasticity, and (iii) inhibitory plasticity (spike timing-dependent plasticity) within CA1. Hebbian updates depended on coincident CA3 activity and basal depolarization, whereas burst-dependent learning required coincident basal and apical input. Silencing ECIII abolished bursts and prevented CA1 remapping in the novel environment, leading instead to reinstatement of the familiar map via SC-driven pattern completion; Hebbian plasticity alone therefore stabilizes familiar CA3→CA1 coupling (F2vsN1, Opsin). In contrast, novelty-driven bursting promoted global CA1 remapping (F2vsN1, Control). Further, inhibitory plasticity was required to stabilize newly acquired maps (N1vsN2).

Together, these results support a dual-input model in which coincidence detection of ECIII and CA3 inputs triggers burst-mediated plasticity able to initiate CA1 remapping, while inhibitory plasticity ensures long-term stability of newly formed spatial maps.

## Discussion

We demonstrate that direct entorhinal input to CA1 is essential for generating a unique CA1 spatial map of a novel environment but has no effect on a familiar map. When ECIII input was inhibited, CA1 failed to undergo global remapping, despite robust CA3 remapping, instead forming a hybrid map combining familiar and novel environment representations. These findings support longstanding models in which CA1 pyramidal neurons act as *mismatch detectors*, comparing ECIII-derived ongoing sensory input with CA3-retrieved representations, or predictions, to detect novelty and continuously reshape local cell assemblies ^17–19,23^.

Specific spatiotemporal ECIII-CA1 input modulates synaptic plasticity at the CA3-CA1 synapse and ECIII-mediated dendritic depolarization as well as associated somatic plateaus and burst firing can rapidly induce new place fields ^39,40,47,50–55^. Thus, direct pathway inhibition likely limits CA1 plasticity, biasing activity towards previously potentiated synapses. Yet this alone cannot explain the reactivation of familiar spatial tuning in the novel environment, as CA3, the remaining input to CA1, exhibited robust remapping. Our model reveals that minimal overlap in CA3 representations, combined with extensive convergence of CA3 inputs onto CA1 neurons ^56^, is sufficient to reinstate familiar CA1 spatial firing patterns when ECIII input is absent in the novel environment.

The persistence of the hybrid map upon re-exposure to the novel environment despite intact ECIII input, indicates that CA1 plasticity is temporally gated, preventing effective reorganisation of spatial tuning once the *novelty state* subsides. The modulation of such a novelty state could be attributed to different mechanisms. Indeed, novelty is associated with dendritic spikes, dendritic disinhibition and enhanced neuromodulatory drive, all of which decline with familiarisation ^17,52,57–61^. In our model, however, temporal gating emerges from the effects of Hebbian plasticity, which progressively aligns CA3 and CA1 representations, prevents burst firing and eventually stops further map reorganisation.

We propose that CA1 global remapping emerges from the interaction between novelty detection and a transient plasticity-permissive state that enhances the influence of ECIII input. This dual mechanism enables the hippocampus to balance the formation of distinct representations of new experiences with the preservation of existing memories.

### Limitations of the study

Due to light scattering in tissue and the inherent difficulty in manipulating axon terminals ^62^, our silencing of ECIII inputs may have been incomplete. In addition, the laser was interrupted during immobility to prevent detrimental side effects. This may therefore explain the partial blockade of remapping observed in Figure 2.

Inhibitory opsin expression in our transgenic mouse was homogenous across medial and lateral EC (Figure 1A) ^41,43^, preventing us from isolating specific contributions of these entorhinal subregions. Prior work observed that CA1 can exhibit global remapping when medial EC is lesioned ^63^, implying that lateral EC conveys substantial spatial information ^64^.

Our tetrode recordings were restricted to CA1 *stratum pyramidale*, and sampled primarily putative pyramidal excitatory neurons, limiting our ability to assess how ECIII input inhibition affects interneuron networks during remapping.

## ACKNOWLEDGEMENTS

We thank Q. Zhang and C. Schrick for *in situ* hybridization and the Kentros group for helpful discussions through the project. This work was supported by European Union’s Horizon 2020 research and innovation program (MGate, grant agreement no. 765549; O.C., M.G., L.D., C.K. and F.P.B.), the European Research Council (ERC) Advanced Grant “REPLAY-DMN” (grant agreement no. 833964; F.P.B.), and European Union’s Horizon 2020 Research and Innovation Program Grant “Brownian Reactivation” (grant agreement no. 840704; F.D., F.S.), FRIPRO ToppForsk grant Enhanced Transgenics (90096000) of the Research Council of Norway (O.C., L.D. and C.K), the Kavli Foundation (O.C., L.D. and C.K), the Centre of Excellence scheme of the Research Council of Norway-Centre for Biology of Memory and Centre for Neural Computation (O.C., L.D. and C.K), The Egil and Pauline Braathen and Fred Kavli Centre for Cortical Microcircuits (O.C., L.D. and C.K), and the National Infrastructure scheme of the Research Council of Norway-NORBRAIN (O.C., L.D. and C.K).

## AUTHOR CONTRIBUTIONS

Conceptualization: O.C., M.G., F.S., F.P.B. and C.K.; Investigation and Data Curation: O.C. and M.G.; Methodology and Software: O.C., M.G., F.D., L.D., F.S., F.P.B and C.K.; Resources: F.S., F.P.B. and C.K.; Formal Analysis and Visualization: O.C., M.G., F.D. and L.D.; Writing – original draft: O.C., M.G. and C.K.; Writing – review and editing: O.C., M.G., F.D., L.D., F.S., F.P.B. and C.K.; Supervision and Funding Acquisition: F.S., F.P.B, C.K.

## COMPETING FINANCIAL INTERESTS

The authors declare no competing interests.

## Methods

### Subjects

11 male mice (16-31 weeks of age) were used in this study. 8 mice were implanted with a Hybrid Drive ^65^ and 3 mice with a Shuttle Drive ^66^. Animals were kept on a 12-h light-dark cycle and individually housed in an enriched environment. Water and food were available *ad libitum*. All procedures and experiments were performed in accordance with the Norwegian Animal Welfare Act and the European Convention for the Protection of Vertebrate Animals used for Experimental and Other Scientific Purposes. Protocols were approved by the Norwegian Food and Safety Authority (FOTS ID 17898).

### Transgenic mouse lines

The transgenic mouse lines used in this study are the same as those previously described in Guardamagna *et al*. ^43^. To obtain specific transgene expression in entorhinal cortex layer III (ECIII) cells we used the ECLIII-tTA driver line developed using the Enhancer Driven Gene Expression method ^41^. The ECLIII-tTA driver line was crossed to a tTA-dependent JAWS-tetO line to allow optical inhibition of the direct entorhinal pathway to the hippocampal area CA1.

### Surgery

All surgical procedures were performed under anesthesia with isoflurane (5% induction, 1-2% maintenance), plus a subcutaneous injection of analgesics (Metacam, 5mg/kg and Temgesic, 0.1mg/kg) and a local anesthesia (Marcain, 1mg/kg) prior to surgery. For electrophysiological tetrode recordings, 7 mice were implanted with a Hybrid Drive ^65^ and 3 animals were implanted with a Shuttle Drive ^66^. The Hybrid Drive was equipped with 13 while the Shuttle Drive contained 16 independently movable tetrodes targeting hippocampal area CA1 *stratum pyramidale*. Anesthetised animals were secured in a stereotaxic frame, and a rectangle craniotomy was performed above the right hippocampus at coordinates −1.2mm anteroposterior and 0.6mm lateral from bregma for the top left corner and −2.3mm anteroposterior and 2.1 lateral from bregma for the bottom right corner. Following the implantation of the drive, sterile Vaseline was used to cover the exposed parts of the tetrodes and dental cement (Superbond C&B, Sun Medical, Japan) plus 3 stainless steel M0.8 anchor screws were used to secure the drive to the skull. Mice recovered for at least 7 days before experiments began.

For optogenetic silencing of ECIII projections to CA1, an optic fiber was incorporated in both the Hybrid and Shuttle Drive (core diameter: 100µm, numerical aperture: 0.37, Doric Lenses) and implanted at a depth of 1mm.

To assess the effect of direct ECIII inhibition on CA1 dynamics we compared transgenic animals expressing the inhibitory opsin JAWS ^42^ (*n*=7 mice – Opsin group) with control littermates that did not express the opsin (*n*=4 mice – Control group). Simultaneous hippocampal CA1 and CA3 electrophysiological tetrode recordings were performed in 2 Opsin mice by targeting 8 tetrodes of the Shuttle Drive to each subregion while the optic fiber implant, as for all animals, targeted CA1.

### Electrophysiological and behavioural data collection

Extracellular signals were acquired using an Open Ephys Acquisition Board ^67^. Signals were referenced to ground, bandpass filtered at 1 and 7500Hz, amplified, multiplexed and digitized at 30kHz on the headstage (for the Hybrid Drive: RHD2132, Intan Technologies, USA; for the Shuttle Drive: RHD2164, Intan Technologies, USA). Digital signals were transmitted over 2 custom 12-wire cables (CZ 1187, Cooner Wire, USA). Waveform extraction and automated clustering were performed using Dataman (https://github.com/wonkoderverstaendige/dataman). Clustered units were manually curated using the MClust spike sorting toolbox. Behavioural data was acquired using a video camera (The Imaging Source, DMK 37BUX273) mounted above the recording environments. Extracellular signals from each tetrode were assessed during rest sessions in the animal’s home cage placed in the recording room. Tetrodes were gradually lowered (45/60µm per day) until clear physiological markers of CA1 and CA3 *stratum pyramidale* were observed, e.g. sharp wave ripple complexed during sleep and theta during exploratory behaviour. On average the target location was reached within 10 days.

### Behavioural paradigm

All mice were familiarised to a standard open field box (50×50×50cm), environment A, with 3 white walls and one black wall. The environment was considered familiar after a minimum of 6 exposures of 15-20 minutes over the course of 2 days prior to the beginning of the experiment. The novel open field box (50×50×35cm), environment B, had 3 black walls and one white wall, a textured floor (untextured in the familiar box), distinct lighting arrangements and was scented with 1% acetic acid (no scent in the familiar box).

On the experimental day, mice underwent 5 consecutive sessions of 15-20 minutes, each separated by a 5-minute break in their homecage. Mice were first exposed to the familiar environment (environment A; session F1), then reintroduced to the familiar environment (environment A; session F2) with optogenetic inhibition of ECIII projection terminals in CA1. Next, mice were placed in the novel environment (environment B; session N1) with optogenetic inhibition of ECIII projection terminals in CA1 before being re-exposed to both familiar (environment A; session F3) and novel (environment B; session N2) environments without optogenetic inhibition. Running in each environment was motivated by scattering chocolate crumbs uniformly across the box.

### Optogenetic manipulation of ECIII terminals

Optogenetic silencing of ECIII projections to s*tratum lacunosum-moleculare* of the CA1 subregion of the dorsal hippocampus was performed by delivering continuous red light (640nm, 20mW input power; 16mW output power) by a diode laser (Coherent Obis LX 1185054). To avoid excessive light stimulation and its detrimental effects ^55^, we confined the length of light delivery with two conditions: (1) the laser was turned on during periods of exploration and turned off during periods of immobility, (2) if exploration periods exceeded 60 seconds, the laser was turned off for minimum 5 seconds before being turned on again. To evaluate the extent of inhibition across the arena we computed a stimulation ratio defined as the time spent with the laser on divided by the total time spent within each spatial bin (Figure S1B and S1C). Optogenetic inhibition covered equally well the arena in both the Opsin and Control groups (Figure S1B and S1C) and lasted on average over 85% of the duration of each session (F2 and N1), with no significant difference between Opsin (85.5±1.7%, mean±SEM) and Control group animals (91.4±1.1%, mean±SEM) (Figure S1C). To assess the involvement of the ECIII input in CA1 spatial coding, we combined this optogenetic manipulation with population recordings of CA1 pyramidal cells (Opsin group: 31.7±5.7 cells; Control group: 23.5±6.3 cells; mean±SEM; Figure 1B and Figure S1A).

### Histology

Histological procedures were performed as previously described in Guardamagna et al. ^43^. At the conclusion of the experiment, mice were euthanised with an overdose of pentobarbital and transcardially perfused with RNase free 4% paraformaldehyde dissolved in PBS. The brains were extracted and stored in RNase free 4% paraformaldehyde dissolved in PBS for 24 hours, then transferred to RNase free 30% sucrose for 48 hours. The brains were sliced into 30 µm thick sections using a Cryostat (Fisher Scientific, Cryostar NX70). The right hemisphere was used to identify recording sites and the position of the optic fiber. The brain was sectioned coronally, and the tissue was stained with Cresyl violet. Slides were coverslipped and brightfield images were taken at a magnitude of 5x using a scanner (Zeiss Axio Scan.Z1, Germany). The left hemisphere was used to validate transgene expression. The brain was sectioned sagittally and the tissue was stained using both fluorescent in situ hybridisation and immunohistochemistry as described in Guardamagna et al. ^43^.

### Data analysis

#### Place cell identification and classification

All recordings were velocity filtered for periods when mice were running over 2cm/s. Putative excitatory pyramidal cells were separated from putative interneurons using auto-correlograms, firings rates and waveform properties. A unit was classified as a pyramidal cell if the mean firing rate was < 8Hz and the average first moment of the auto-correlogram occurred under 8ms. Putative excitatory neurons were then subjected to further place cell analysis. For each session, rate maps were generated by binning position data in 2×2cm bins and dividing the spike count by the time spent in each bin, then applying a Gaussian kernel with a standard deviation of 2 bins. Place cells were defined using the following criteria: (1) mean firing rate 0.3Hz; (2) peak firing rate 1 Hz; (3) within session spatial stability, i.e. Pearson’s correlation between first and second half of the session >0.5; (4) at least one identified place field, defined as areas with at least 18 contiguous pixels where the firing rate exceeded 20% of the peak rate; (5) Skaggs information content (bits/spike) ^68^ significantly higher than shuffled data. Shuffling was performed by randomly shuffling cell spike times 1000 times per cell, generating a distribution of values. If a cell’s information content exceeded the 95^th^ percentile of the shuffled distribution, it was considered significant. Skaggs information content:

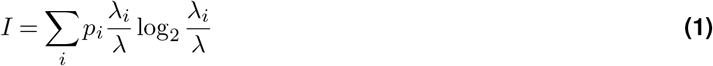

where *p*_*i*_ is the probability of the mouse being in the *i*-th bin, *λ* is the mean firing rate, *λ*_*i*_ is the mean firing rate in the *i*-th bin. Only cells defined as place cells in at least one session were used for further analysis.

#### Remapping analysis

To evaluate the reorganisation of spatial tuning across sessions following ECIII input inhibition, we compared Pearson’s correlation coefficients of place cells between distinct sessions. Only cells with a firing rate >0.1Hz in at least one of the two sessions compared were included. To determine remapping kinetics, all sessions were split into one-minute epochs, and for all cells, rate maps for each epoch were computed. Then, to assess the speed of decorrelation upon entrance to the novel environment, Pearson’s correlation coefficients were computed between rate maps from each one-minute epoch and rate maps generated from the entire familiar environment session without ECIII input inhibition (F1). Likewise, to assess the speed of emergence of the novel environment map during ECIII input inhibition, Pearson’s correlation coefficients were computed between rate maps from each one-minute epoch and rate maps generated from the entire first novel environment session (N1). Comparisons were only computed if the animal covered over 10% of the environment in the one-minute epoch. The displacement or shift of place fields between sessions was quantified as the Euclidean distance of a cell’s place field center-of-mass (COM).

To compare spatial maps at the population level across sessions, all rate maps from a single session were stacked into a three-dimensional matrix, where the *x-y* axes corresponded to the arena position and the *z*-axis to the cell identity. Population vector (PV) correlations are Pearson’s correlations for each *x-y* location, i.e. spatial bin, between two different sessions, defining a correlation score per bin. Alternatively, the same process was applied to rate maps of individual animals separately, then correlations scores from all spatial bins were averaged per animal. Only animals with over 10 simultaneously recorded place cells were used for this analysis. To visualize the degree of remapping for simultaneously recorded place cells across sessions, we computed spatial cross-correlograms ^30^. For each session, rate maps were stacked into a three-dimensional matrix, as described above. To calculate the cross-correlation dot product between PVs of different sessions, one stack was shifted relative to the other in 2cm (same size used to discretize spatial dimensions for rate maps) increments along the *x* and *y* axes. Cross-correlograms are displayed as a two-dimensional 49×49 matrix of 2cm bins.

#### Rate change

The mean firing rate of individual place cells was calculated by dividing the total number of spikes by the duration of the recording session. To assess changes in firing rates across sessions, a difference score was calculated as following:

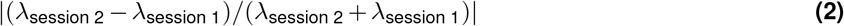

where λ is the mean firing rate for a given session.

#### Overlap

To assess the *overlap* ^13^ of active cells between sessions, we computed for each cell the ratio between mean firing rate in the session with the least active by the mean firing rate in the session the most active.

#### Burst firing

Bursts were defined as spike trains with inter-spike intervals <9ms. To quantify the effect of ECIII input inhibition on burst firing in CA1 place cells, we compared the change in burst probability between sessions. Burst probability:

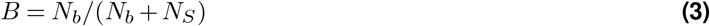

where *N*_*b*_ is the total number of bursts and *N*_*s*_ the number of single spikes for a given session. Change in burst probability was calculated as a difference score:

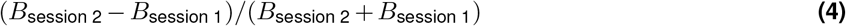

To characterize whether the probability of a place cell to burst was homogeneous throughout the session or was enhanced during a specific period, we calculated the burst probability of individual place cells as a function of time spent in its place field (or eventual place field location) for one-minute epochs. We restricted this analysis to time within the place field to correct for irregular exploration throughout the environment.

### Spiking neural network model of the hippocampal circuit

To investigate the neural dynamics underlying remapping in CA1, we implemented a computational model using a network of two-compartment Leaky Integrate-and-Fire (LIF) neurons. The model was designed to reproduce key experimental findings and to explore the contributions of inhibitory plasticity to spatial learning and memory

#### Model specifics

The pyramidal cells in CA1 were modeled as a two-compartment LIF neuron, with an apical (A) and basal (B) compartment. Moreover, two types of single-compartment interneurons are being modeled: one which specifically inhibits the apical compartment (“SOM”) of the pyramidal cells, and one which inhibits the basal compartment (“PV”). For a visualization of the architecture see Figure 4D. Neuron *i* in population *k* adheres to the following dynamics:

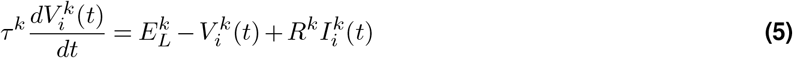

where *k* ∈ {A, B, SST, PV} is the type of cell or compartment, *τ*^*k*^ the time constant, 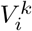 the voltage, 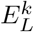 the leak reversal potential and reset voltage, *R*^*k*^ the resistance, and 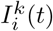 the input.

The input to each type differs as follows:

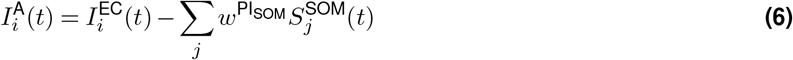

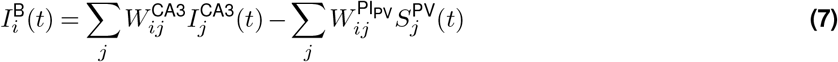

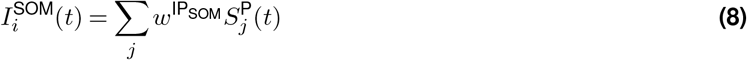

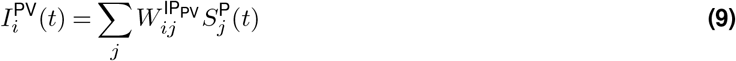

Where 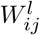 are connection matrices for *l* ∈ {PI_PV_, CA3, IP_PV_} while 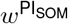 and 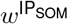 are constant connection weights. 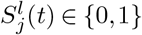 is a binary variable indicating spikes at each simulation step for *l* ∈{SST, PV, P}. *P* refers to the output of a pyramidal cell, determined from the state of both compartments.

Additionally, to these dynamics, there are several reset conditions. For each of the interneuron types and for each of the compartments of the pyramidal cells, there is a threshold 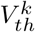. For the interneurons, if this threshold is reached, the voltage will be reset, 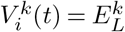 and the neuron will spike, i.e. 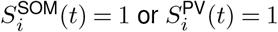.

However, the pyramidal cells can be in one out of three states depending on which thresholds are reached: the neuron can be silent, spiking, or bursting. Bursts are represented by a binary variabl 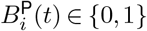. If only the basal compartment reaches its threshold, the neuron spikes, 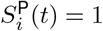. If both the basal and the apical compartments reach their respective thresholds, it bursts 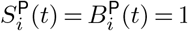. In all other cases, the neuron is silent, 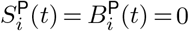. Hence, in our model, a burst is explicitly defined as a simultaneous crossing of the apical and the basal threshold. Any activity of the neuron will be referred to as an event while the case 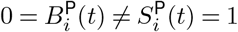 will be referred to as spike and the case 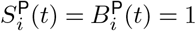 compartment are reset.

will be referred to as burst. After every event, both the apical and the basal For all simulations, all connections were all-to-all such that each CA1 unit received input from all CA3 cells and was connected to all inhibitory neurons which, in turn, were again connected to all CA1 cells. The initial condition of all cells and compartments was chosen as 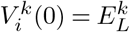. The dynamics were simulated for each neuron/compartment type sequentially using a second-order Runge-Kutta solver in Python with a simulation timestep Δ*t* (see Table S2).

#### Excitatory plasticity

The connection matrix representing the Schaffer Collaterals, i.e. 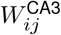 is subject to plasticity. The weights are initialized randomly and drawn from a uniform distribution, 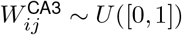. The plasticity then depends both on top-down input from the ECIII as well as on Hebbian plasticity (see Figure 4D),

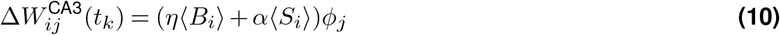

where *i* denotes the postsynaptic neuron in CA1, *j* the presynaptic neuron in CA3 and ⟨·⟩ denotes a time average of spikes *S*_*i*_ or bursts *B*_*i*_ over a specified plasticity time window, Δ*t*_plast_. *ϕ*_*j*_ is a trace of activity of neuron *j* in CA3, i.e. the mean activity over the plasticity time window. While *η* controls the strength of the burst dependent plasticity, *α* controls the strength of Hebbian plasticity. Since we are modelling excitatory cells in CA3, Dale’s Law is ensured by clipping the weights to a minimum of zero after each update,

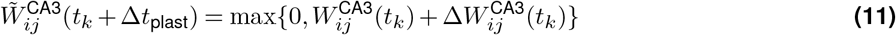

Moreover, the weights are normalized so that, at all times, the total incoming weight to each cell remains constant. This is achieved by normalizing over all presynaptic indices *j* for a given postsynaptic index *i*, i.e.,

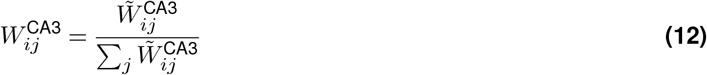

This ensures that each CA1 neuron’s total incoming Schaffer collateral weight remains constant.

#### Inhibitory plasticity

Additionally, we implemented a spike timing-dependent plasticity (STDP) rule for the PV-like cells in the simulation which targets the basal compartment of the pyramidal cells. In the experiments where the inhibitory weights were not subject to plasticity, they were instead constant, 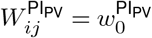and 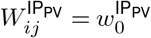 for all *i, j*. For all simulations with inhibitory plasticity, the inhibitory weights were initialized from a uniform distribution such that 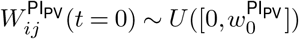 and 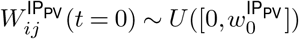. The STDP rule depends on the momentary spiking behaviour of presynaptic neuron *j* and postsynaptic neuron *i* and a synaptic trace defined for both neurons *k* ∈ {*i, j*} as

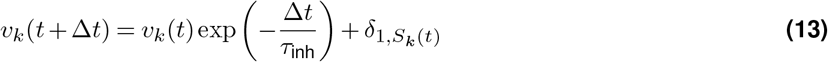

where *S*_*k*_(*t*) ∈ {0, 1} is a binary variable indicating spiking behaviour of neuron *k* and *δ* is the Kronecker delta. The *Excitatory* → *Inhibitory* weights are adapted with the following STDP rule:

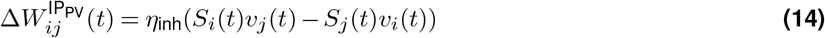

The *Inhibitory* → *Excitatory* weights, however, are adapted with a symmetric STDP rule combined with a constant depression term:

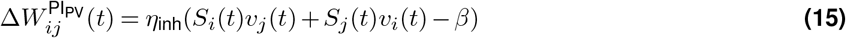

where *β* is the depression constant. As in the excitatory case, Dale’s Law is enforced by keeping zero as a lower bound:

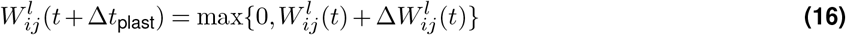

for *l* ∈ {IP_PV_,PI_PV_}

#### Trajectory simulations

To generate the activity patterns of CA3 cells and the top-down input from the ECIII, an animal was simulated to move through a two-dimensional box. To avoid edge effects, the environment was modeled as a torus, i.e., the edges are connected. Below, *t* denotes the simulation time, Δ*t* the simulation time step, and *t*_*N*_ the total simulation duration.

#### Trajectory

The animal moves within a square environment of side length *L*. Its position is **x**(*t*) = (*x*(*t*), *y*(*t*)), starting at the centre **x**(0) = (*L/*2, *L/*2). The scalar speed *v*(*t*) has an initial value *v*(0) = *v*_0_, and the initial movement direction *ϕ*(0) is drawn from a uniform distribution, *ϕ*(0) ~ *U* ([0, 2*π*)). At each simulation step, the direction and speed evolve as:

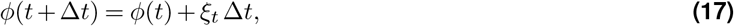

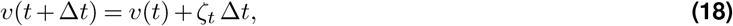

where *ξ*_*t*_ ~ N (0, *σ*_*ϕ*_) and *ζ*_*t*_ ~ N (0, *σ*_*v*_). The displacements in *x* and *y* direction are then simply given by:

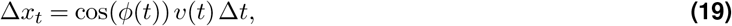

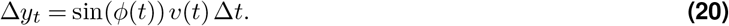

Periodic (toroidal) boundary conditions are applied independently to both coordinates:

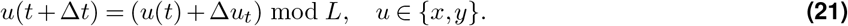

This ensures that crossing one boundary leads to re-entry on the opposite side.

#### Learning Problem

Activity in CA3 and input from the EC were modeled as Gaussian functions of the animal’s position:

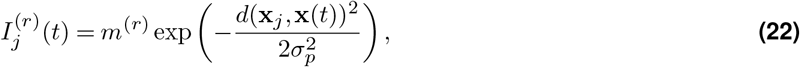

where *r* ∈{EC, CA3} denotes the input source, *m*^(*r*)^ is the maximum input strength for population *r*, **x**_*j*_ denote the place field centres and **x**(*t*) the position of the animal at time point *t*. We define **x**_*j*_ = (*x*_*j*_, *y*_*j*_) and **x**(*t*) = (*x*(*t*), *y*(*t*)). *d*(·, ·) denotes the distance metric, which, since the animal is modeled on a torus, is given by the wrapped (torus) distance. For each coordinate *u* ∈ {*x, y*}, the wrapped distance is defined as

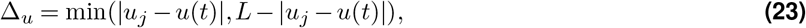

where *L* is the edge length of the square environment. The full 2D wrapped distance is then

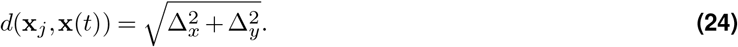

CA3 activity is modeled per CA3 unit in terms of firing rates, whereas EC input is assumed to have already been transformed into place field activity that directly enters the apical compartment of each pyramidal cell as can be seen in the equations above. The learning problem therefore consists of associating CA3 activity with the corresponding top-down signal from the EC. Place field centres *x*_*j*_ are evenly spaced within the square and randomly permuted. Two distinct environments are simulated by randomly reshuffling the place field centres.

#### Experimental paradigm

To study the remapping behaviour of the model, we used the same paradigm as in the experimental part. To simulate the two experimental groups, we proceeded as follows: For the *experimental group*, the top-down entorhinal cortex (EC) input to CA1 is silenced during stages F2 and N1, while for the *control group* the EC input is present during all stages.

To simulate the familiar and the novel environments respectively, two separate maps were simulated as discussed above. To simulate a certain correlation between different environments, maps in CA3 from the novel environment were created as a linear combination of two Gaussians with different centres such that

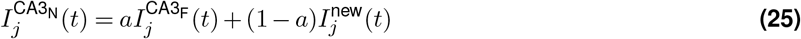

Maps for the top-down ECIII input, however, were uncorrelated and created as discussed above. As for the experimental behavioural paradigm, we sequentially simulated five different stages (or sessions) for two groups mimicking the Opsin and Control group conditions.

In session F1, a run was simulated in the familiar environment, with both groups receiving the CA3 and ECIII inputs. In session F2, the familiar environment was presented again; to simulate optogenetic silencing of the direct entorhinal input to CA1, in the Opsin group condition top-down ECIII input was removed, while in the Control group condition the ECIII input was as in session F1. Both groups received the same CA3 input as in session F1.

In session N1, the novel environment was presented for the first time, with ECIII input silenced in the Opsin group condition and new EC maps presented in the Control group condition; both groups received a mixed CA3 map. In session F3, the familiar environment was presented again, and in session N2, the novel environment was repeated, with inputs presented as in sessions F1 and N1, respectively.

#### Model statistical analysis

To assess remapping between experimental stages, stage pairs were compared using the following measures of association: (1) Pearson’s correlation and (2) matrix cross-correlation. Spike trains were temporally smoothed, yielding activity maps of dimension *n*_pos_ × *n*_cells_, where *n*_pos_ is the number of spatial bins and *n*_cells_ the number of neurons. Measures (1) and (2) were computed as described above for experimental data.

#### Simulations

The model was run sequentially through the different experimental stages. For each experimental stage and group, we computed an activation map by binning the environment into 64 spatial bins over the full simulation. The average firing rate in each bin was then calculated using a rectangular kernel. We then computed PV and spatial correlations as well as cross correlograms between the following pairs of activation maps: F1 vs F2, F2 vs N1, N1 vs N2. Then, to study the importance of the inhibitory plasticity to reproduce the experimental results, the experimental paradigm was repeated while silencing the inhibitory STDP. While all other parameters were kept at the same values (Table S2), the experimental paradigm was repeated, setting *η*_inh_ = 0. Then, again, the same pairwise comparisons between experimental stages as above, spatial and PV correlations as well as cross-correlograms were computed.

### Statistics

To obtain consistent and homogenous opsin expression across animals, we used a transgenic mouse. Following genotyping, double-positive animals (expressing the opsin) were assigned to the Opsin group, and single-positive littermates (that did not express the opsin) were assigned to the Control group. Data analyses were performed using custom MATLAB (MathWorks) or Python scripts. Nonparametric tests were used when assumptions for parametric tests were not met, and median values were reported. All statistical tests were performed using standard built-in MATLAB functions. Error is reported as standard error of the mean (SEM). \**p*<0.05; \*\**p*<0.01; \*\*\**p*<0.001.

## Data availability

Data will be publicly available following publication.

## Supplementary Figures

**Figure S1.**
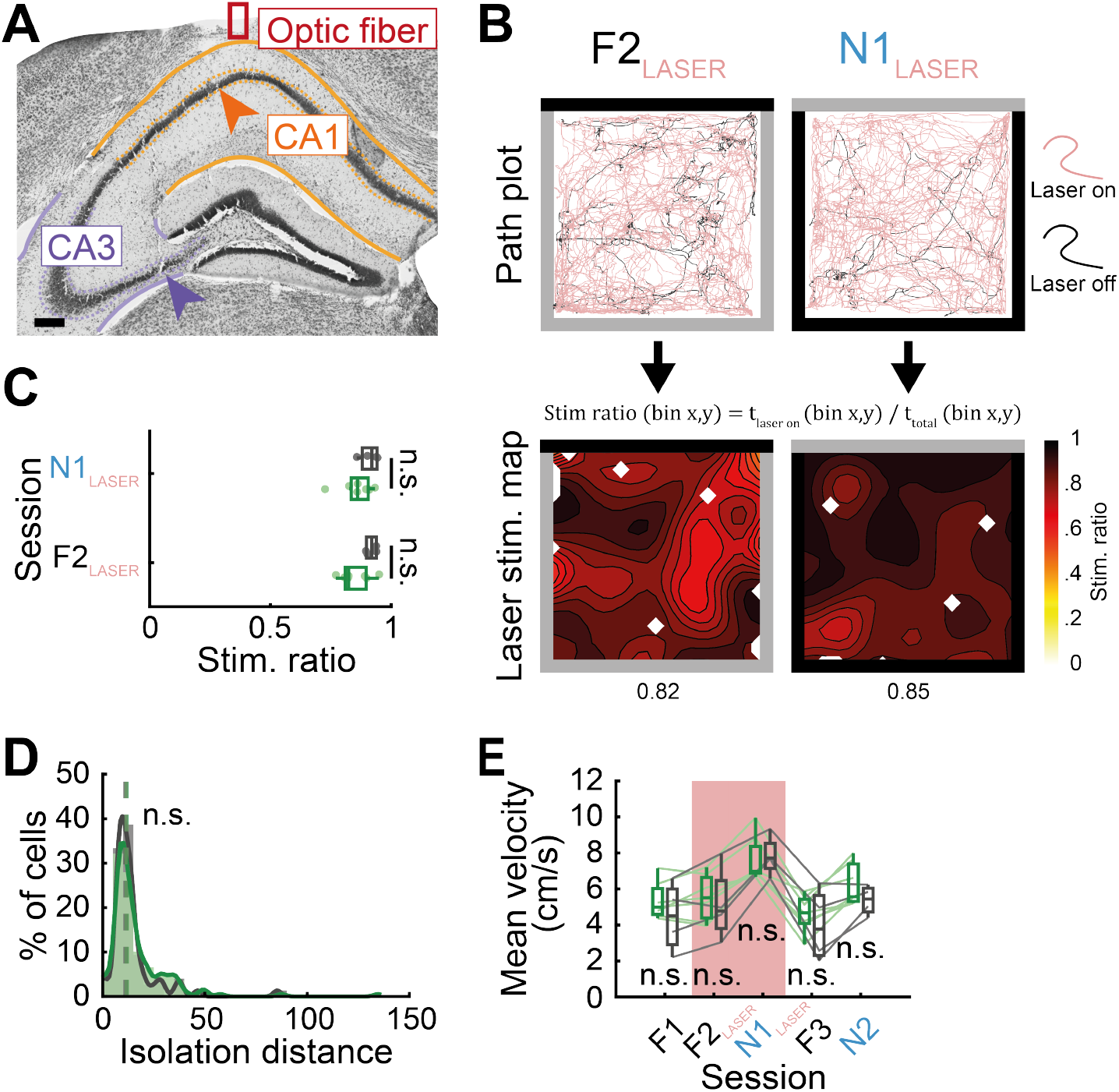
Hippocampal recordings and optogenetic manipulation in freely exploring mice. **(A)** Nissl-stained coronal section showing an example optic fiber tract (red rectangle) and tetrode recording sites in CA1 (orange arrow) and in CA3 (purple arrow). Scale bar: 200µm. **(B)** Optogenetic stimulation strategy. Top: example path plots during familiar and novel environments sessions with laser stimulation. Red and black paths represent periods with the laser on and off, respectively. To quantify stimulation coverage, we computed a stimulation ratio for each spatial bin (2cmx2cm) as the fraction of the time with laser on relative to the total time. Bottom: contour plots of stimulation ratios across familiar and novel environments. Values below each plot represents arena-wide mean stimulation ratios. **(C)** Average stimulation ratios for both Opsin (*n*=7) and Control (*n*=4) animals. F2: *p*=0.0936; N1: *p*=0.1843, permutation test. **(D)** Distribution of unit isolation distances for all recorded units from the Opsin (*N*=168) and Control (*N*=75) groups. *p*=0.3286, Wilcoxon rank-sum test. **(E)** Mean running velocity across all sessions for all Opsin (*n*=7) and Control (*n*=4) animals. F1: *p*=0.3413; F2: *p*=0.6767; N1: *p*=0.7221; F3: *p*=0.4713; N2: *p*=0.2296, permutation test. Box plots show median and interquartile range. Dashed vertical lines overlaying histograms show medians. Green, Opsin; gray, Control.

**Figure S2.**
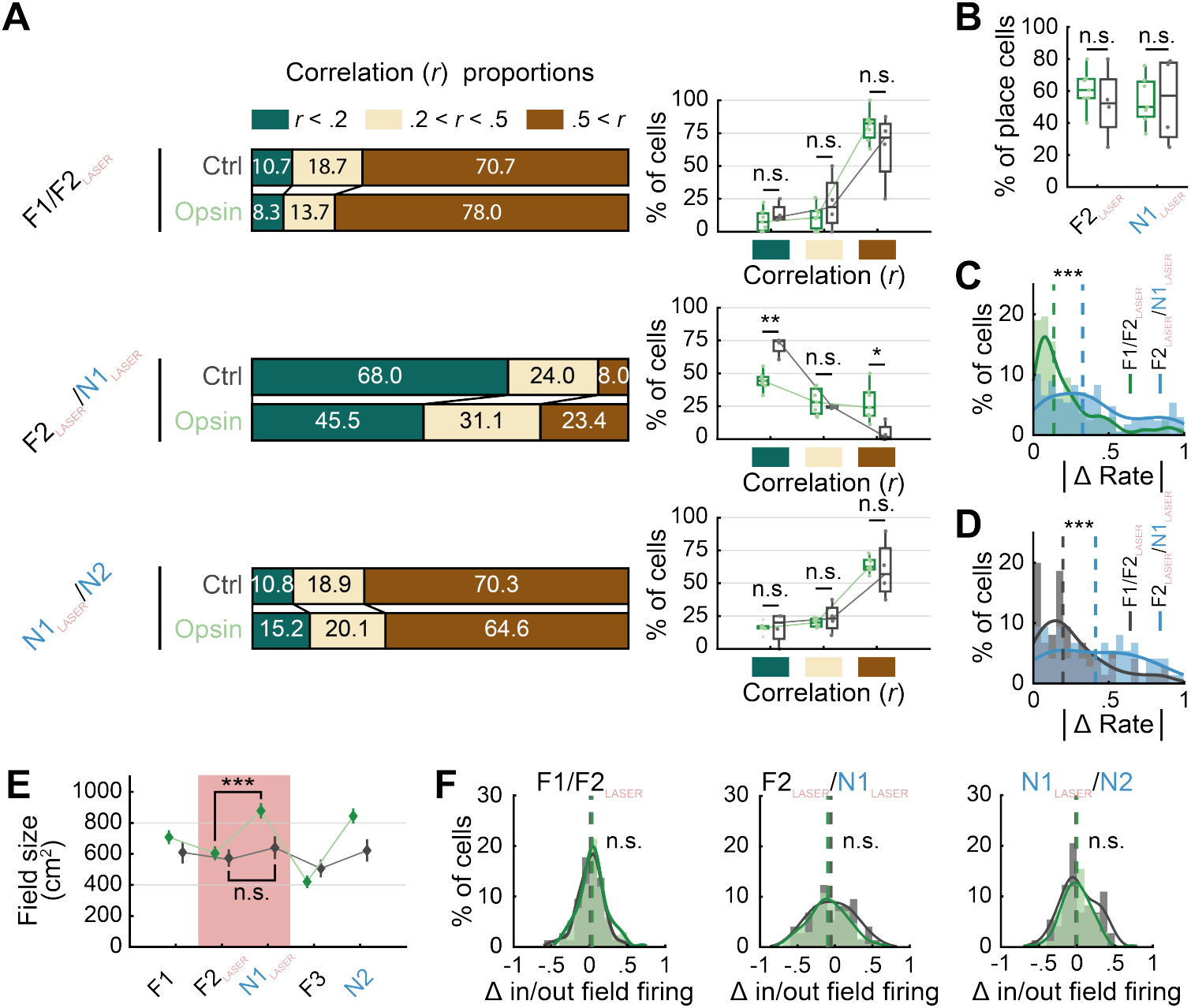
The effect of ECIII input inhibition on place cell firing properties in a familiar and novel environment. **(A)** Left: proportions of Opsin (*N*=168) and Control (*N*=75) place cells with low (green, *r* < 0.2), intermediate (beige, 0.2 < *r* < 0.5) and high (brown, *r* > 0.5) Pearson correlation coefficients between sessions F1 and F2 (top), F2 and N1 (middle) and, N1 and N2 (bottom). Right: average proportions per animal (Opsin, *n*=7; Control, *n*=4). Between sessions F1 and F2 (top). *r* < 0.2, *p*=0.2960; 0.2 < *r* < 0.5, *p*=0.2658; *r* > 0.5, *p*=0.1963. Between sessions F2 and N1 (middle). *r* < 0.2, \*\**p*=0.0030; 0.2 < *r* < 0.5, *p*=0.4260; *r* > 0.5, \**p*=0.0181. Between sessions N1 and N2 (top). *r* < 0.2, *p*=0.9698; 0.2 < *r* < 0.5, *p*=0.4804; *r* > 0.5, *p*=0.6798. Permutation test. **(B)** Fraction of place cells for Opsin (*n*=7) and Control (*n*=4) animals during familiar (F2) and novel (N1) sessions with ECIII input inhibition. For session F2, *p*=0.4290; for session N1, *p*=0.9799. Permutation test. **(C)** Distribution of place cell absolute firing rate change for the Opsin group (*N*=168) between familiar sessions without (F1) and with (F2) ECIII input inhibition (green) and between familiar (F2) and novel (N1) sessions with ECIII input inhibition (blue). \*\*\**p*=5.1×10^−13^, Wilcoxon signed-rank test. **(D)** Same as **(C)** for the Control group (*N*=75). \*\*\**p*=1.7×10^−4^, Wilcoxon signed-rank test. **(E)** Average place field size (median±SEM) for Opsin and Control groups across sessions. Between sessions F2 and N1: Opsin group (F2, *N*=140 place fields; N1, *N*=132 place fields): \*\*\**p*=3.3×10^−4^; Control group (F2, *N*=60 place fields; N1, *N*=64 place fields): *p*=0.0593, Wilcoxon signed-rank test. **(F)** Distribution of in-field versus out-of-field firing rate ratio change. Left: place fields between sessions F1 and F2 (Opsin, *N*=123; Control, *N*=54), *p*=0.3102, Wilcoxon rank-sum test. Place fields between sessions F2 and N1 (Opsin, *N*=106; Control, *N*=53), *p*=0.3449, Wilcoxon rank-sum test. Place fields between sessions N1 and N2 (Opsin, *N*=114; Control, *N*=58), *p*=0.5098, Wilcoxon rank-sum test. Box plots show median and interquartile range. Dashed vertical lines overlaying histograms show medians. Green, Opsin group; gray, Control group.

**Figure S3.**
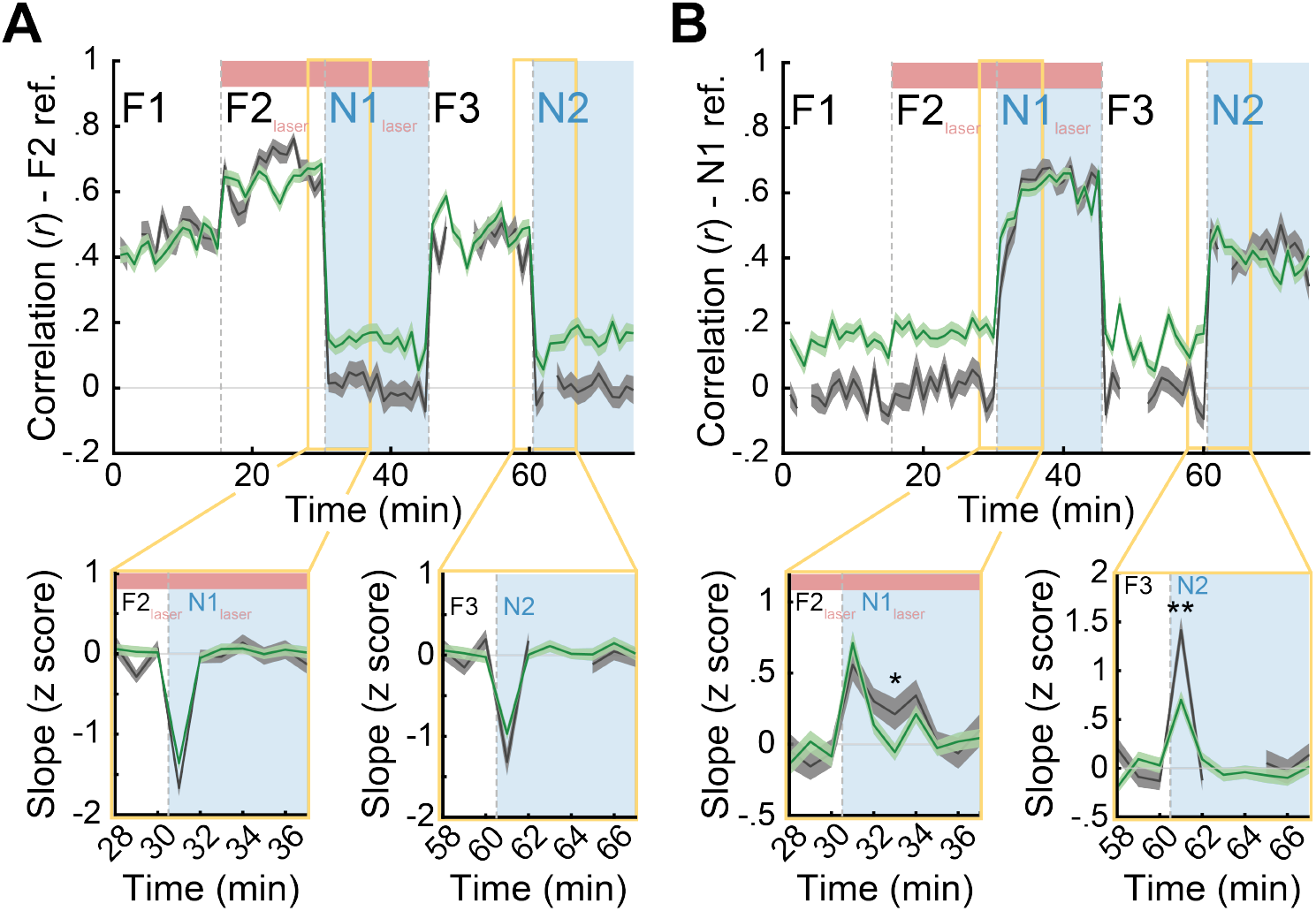
Remapping kinetics. **(A-B)** Top: Pearson’s correlation between 1-minute block rate maps and **(A)** the entire familiar session rate map (F2; reference, ref), and **(B)** the entire first novel session rate map (N1; reference, ref) (Opsin, *N*=168; Control, *N*=75). Bottom: expanded view of the z-scored derivative of the Pearson’s correlation shown above, indicating the rate of representational change between consecutive minutes for transitions between familiar and novel environment sessions. All panels show medians±SEM. See Table S1 for statistics. Green, Opsin group; gray, Control group.

## Supplementary Tables

**Table S1.** Statistical comparisons of Pearson’s correlation between Opsin and Control groups for each 1-minute block with F2 and N1 as reference sessions. \**p* < 0.05; \*\**p* < 0.01; \*\*\**p* < 0.001

| Minute | Session F2 ref. | Session N1 ref. |
| --- | --- | --- |
|  | *p*-value | *p*-value |
| 1 | 0.7346 | 0.0035** |
| 2 | 0.8395 | 0.0067** |
| 3 | / | / |
| 4 | 0.6029 | 0.0034** |
| 5 | 0.4761 | 0.0015** |
| 6 | 0.9740 | 0.0569 |
| 7 | 0.0615 | 0.0115* |
| 8 | 0.8768 | 0.1497 |
| 9 | 0.4816 | 0.0019** |
| 10 | 0.3430 | 0.0104* |
| 11 | 0.3595 | 0.0018** |
| 12 | 0.4487 | 0.0043** |
| 13 | 0.3859 | 0.1276 |
| 14 | 0.7362 | 0.0001*** |
| 15 | 0.8805 | 0.0074** |
| 16 | 0.4755 | 4.5x10<sup>-5</sup>*** |
| 17 | 0.2683 | 0.0016** |
| 18 | 0.0838 | 0.0008*** |
| 19 | 0.6821 | 0.1919 |
| 20 | 0.9766 | 0.0110* |
| 21 | 0.4954 | 0.0220* |
| 22 | 0.1771 | 0.0038** |
| 23 | 0.0014** | 0.0615 |
| 24 | 0.0002*** | 5.6x10<sup>-5</sup>*** |
| 25 | 0.0396* | 0.0037** |
| 26 | 0.0101* | 0.0062** |
| 27 | 0.4006 | 0.0023** |
| 28 | 0.6622 | 0.0063** |
| 29 | 0.0484* | 0.0001** |
| 30 | 0.2815 | 0.0029** |
| 31 | 0.0065** | 0.0029** |
| 32 | 0.0246* | 0.3608 |
| 33 | 0.0047** | 0.9934 |
| 34 | 0.0099** | 0.6240 |
| 35 | 0.0165* | 0.1146 |
| 36 | 0.0225* | 0.2092 |
| 37 | 0.0006*** | 0.1508 |
| 38 | 0.0103* | 0.7277 |
| 39 | 0.0001*** | 0.7824 |
| 40 | 0.0058** | 0.7353 |
| 41 | 0.0023** | 0.1513 |
| 42 | 0.0350* | 0.2957 |
| 43 | 0.0145* | 0.8758 |
| 44 | 0.5918 | 0.0803 |
| 45 | 0.0125* | 0.4411 |
| 46 | 0.7163 | 0.0001** |
| 47 | 0.0144 | 0.1599 |
| 48 | 0.1046 | 6.3x10<sup>-5</sup>*** |
| 49 | / | / |
| 50 | / | / |
| 51 | 0.2481 | 0.0090** |
| 52 | 0.7457 | 0.0595 |
| 53 | 0.4218 | 0.0812 |
| 54 | 0.7950 | 0.0476* |
| 55 | 0.4594 | 0.0165* |
| 56 | 0.2120 | 0.0070** |
| 57 | 0.7398 | 0.0042** |
| 58 | 0.2842 | 0.3714 |
| 59 | 0.0549 | 0.0013** |
| 60 | 0.6669 | 9.3x10-5*** |
| 61 | 0.0023** | 0.4502 |
| 62 | 0.2820 | 0.2733 |
| 63 | / | / |
| 64 | 0.0297* | 0.3652 |
| 65 | 0.0025** | 0.7725 |
| 66 | 0.0020** | 0.9680 |
| 67 | 0.0002*** | 0.5047 |
| 68 | 0.0007*** | 0.5141 |
| 69 | 0.0064** | 0.3033 |
| 70 | 0.1233 | 0.1045 |
| 71 | 0.0170* | 0.0009*** |
| 72 | 1.7x10-5*** | 0.8949 |
| 73 | 0.0015** | 0.0556 |
| 74 | 0.0019** | 0.2399 |
| 75 | 0.0009*** | 0.0959 |

**Table S2.** Model parameters. These parameter settings were applied consistently across all simulations, unless explicitly stated otherwise.

| LIF Parameters |  | Simulation Parameters |  |
| --- | --- | --- | --- |
| Parameter | Value | Parameter | Value |
| $E_L^k$ | -65 | $\tau_{inh}$ | 0.1 |
| $R^k$ | 10 | $\Delta t_{plast}$ | 1 |
| $V_{th}^k$ | -45 | $\beta$ | 0.01 |
| $w_{SST}^{IP}$ | $5000/N_{CA1}$ | $\sigma_{pf}$ | 6 |
| $w_{SST}^{PI}$ | $75/N_{SST}$ | $\Delta t$ | 0.001 |
| $w_0^{IPV}$ | $750/N_{CA1}$ | $v_0$ | 20 |
| $w_0^{PIV}$ | $100/N_{PV}$ | $a$ | 0.3 |
| $\tau^A$ | 1 | $m^{EC}$ | 800 |
| $\tau^B$ | 0.25 | $m^{CA3}$ | 30 |
| $\tau^{PV}$ | 0.1 | $\eta$ | 15 |
| $\tau^{SST}$ | 0.1 | $\alpha$ | 0.25 |
| | | $\eta_{inh}$ | 150 |
| | | $L$ | 50 |
| | | $t_N$ | 250 |
| | | $N_{CA3}$ | 169 |
| | | $N_{CA1}$ | 289 |
| | | $N_{SST}$ | 30 |
| | | $N_{PV}$ | 30 |

## Notes

### Competing Interest Statement

The authors have declared no competing interest.

## References

1. W. B. Scoville and B. Milner. Loss of recent memory after bilateral hippocampal lesions. Journal of Neurology Neurosurgery and Psychiatry, 20(1):11–21, 1957. doi: DOI 10.1136/jnnp.20.1.11.

2. John O’Keefe and Lynn Nadel. The Hippocampus as a Cognitive Map. Oxford University Press, Oxford, UK, 1978. ISBN 978-0198572060.

3. R. G. M. Morris, P. Garrud, J. N. P. Rawlins, and J. Okeefe. Place navigation impaired in rats with hippocampal-lesions. Nature, 297(5868):681–683, 1982. doi: DOI 10.1038/297681a0.

4. G. Buzsáki and E. I. Moser. Memory, navigation and theta rhythm in the hippocampal-entorhinal system. Nature Neuroscience, 16(2):130–138, 2013. doi: 10.1038/nn.3304.

5. M. E. Hasselmo. Neuronal rebound spiking, resonance frequency and theta cycle skipping may contribute to grid cell firing in medial entorhinal cortex. Philosophical Transactions of the Royal Society B-Biological Sciences, 369(1635), 2014. doi: ARTN20120523 10.1098/rstb.2012.0523.

6. N. T. M. Robinson, L. A. L. Descamps, L. E. Russell, M. O. Buchholz, B. A. Bicknell, G. K. Antonov, J. Y. N. Lau, R. Nutbrown, C. Schmidt-Hieber, and M. Häusser. Targeted activation of hippocampal place cells drives memory-guided spatial behavior (vol 183, pg 1586, 2020). Cell, 183(7):2041–2042, 2020. doi: 10.1016/j.cell.2020.12.010.

7. J. O’Keefe and J. Dostrovsky. The hippocampus as a spatial map. preliminary evidence from unit activity in the freely-moving rat. Brain Research, 34(1):171–175, 1971. doi: 10.1016/0006-8993(71)90358-1.

8. E. C. Tolman. Cognitive maps in rats and men. Psychological Review, 55(4):189–208, 1948. doi: DOI 10.1037/h0061626.

9. Bruce L. McNaughton, Francesco P. Battaglia, Ole Jensen, Edvard I. Moser, and May-Britt Moser. Path integration and the neural basis of the ‘cognitive map’. Nature Reviews Neuroscience, 7(8):663–678, 2006. doi: 10.1038/nrn1932.

10. R. U. Muller and J. L. Kubie. The effects of changes in the environment on the spatial firing of hippocampal complex-spike cells. Journal of Neuroscience, 7(7):1951–1968, 1987.

11. M. A. Wilson and B. L. Mcnaughton. Dynamics of the hippocampal ensemble code for space. Science, 261(5124): 1055–1058, 1993. doi: DOI 10.1126/science.8351520.

12. L. L. Colgin, E. I. Moser, and M. B. Moser. Understanding memory through hippocampal remapping. Trends in Neurosciences, 31(9):469–477, 2008. doi: 10.1016/j.tins.2008.06.008.

13. S. Leutgeb, J. K. Leutgeb, A. Treves, M. B. Moser, and E. I. Moser. Distinct ensemble codes in hippocampal areas ca3 and ca1. Science, 305(5688):1295–1298, 2004. doi: 10.1126/science.1100265.

14. C. B. Alme, C. L. Miao, K. Jezek, A. Treves, E. I. Moser, and M. B. Moser. Place cells in the hippocampus: Eleven maps for eleven rooms. Proceedings of the National Academy of Sciences of the United States of America, 111(52):18428–18435, 2014. doi: 10.1073/pnas.1421056111.

15. Menno P. Witter, Pieterke A. Naber, Theo van Haeften, Willem C. M. Machielsen, Serge A. R. B. Rombouts, Frederik Barkhof, Philip Scheltens, and Fernando H. Lopes da Silva. Cortico-hippocampal communication by way of parallel parahippocampal-subicular pathways. Hippocampus, 10(4):398–410, 2000. doi: 10.1002/1098-1063(2000)10:4<398::AID-HIPO6>3.0.CO;2-K. URL https://doi.org/10.1002/1098-1063(2000)10:4<398::AID-HIPO6>3.0.CO2-K.

16. Cathrin B. Canto, Floris G. Wouterlood, and Menno P. Witter. What does the anatomical organization of the entorhinal cortex tell us? Neural Plasticity, 2008(1):381243, 2008. doi: 10.1155/2008/381243.

17. M. E. Hasselmo and E. Schnell. Laminar selectivity of the cholinergic suppression of synaptic transmission in rat hippocampal region ca1 - computational modeling and brain slice physiology. Journal of Neuroscience, 14(6):3898–3914, 1994.

18. H. Eichenbaum. Declarative memory: Insights from cognitive neurobiology. Annual Review of Psychology, 48:547–572, 1997. doi: DOI 10.1146/annurev.psych.48.1.547.

19. J. E. Lisman and N. A. Otmakhova. Storage, recall, and novelty detection of sequences by the hippocampus: Elaborating on the socratic model to account for normal and aberrant effects of dopamine. Hippocampus, 11(5):551–568, 2001.

20. O. S. Vinogradova. Hippocampus as comparator: Role of the two input and two output systems of the hippocampus in selection and registration of information. Hippocampus, 11(5):578–598, 2001. doi: 10.1002/hipo.1073.

21. Federico Stella and Alessandro Treves. Reorganization of spatial maps in the hippocampal circuit. BMC Neuroscience, 12 (1):P340, 2011. doi: 10.1186/1471-2202-12-S1-P340.

22. T. Geiller, J. B. Priestley, and A. Losonczy. A local circuit-basis for spatial navigation and memory processes in hippocampal area ca1. Current Opinion in Neurobiology, 79, 2023. doi: ARTN102701 10.1016/j.conb.2023.102701.

23. Yusi Chen, Huanqiu Zhang, Mia Cameron, and Terrence Sejnowski. Predictive sequence learning in the hippocampal formation. Neuron, 112(15):2645–2658.e4, 2024. doi: 10.1016/j.neuron.2024.05.024.

24. Houman Davoudi and David J. Foster. Acute silencing of hippocampal ca3 reveals a dominant role in place field responses. Nature Neuroscience, 22(3):337–342, 2019. doi: 10.1038/s41593-018-0321-z.

25. Ruy Gómez-Ocádiz, Massimiliano Trippa, Chun-Lei Zhang, Lorenzo Posani, Simona Cocco, Rémi Monasson, and Christoph Schmidt-Hieber. A synaptic signal for novelty processing in the hippocampus. Nature Communications, 13(1):4122, 2022. doi: 10.1038/s41467-022-31775-6.

26. A. Treves and E. T. Rolls. Computational constraints suggest the need for 2 distinct input systems to the hippocampal ca3-network. Hippocampus, 2(2):189–200, 1992. doi: DOI 10.1002/hipo.450020209.

27. M. Tsodyks and T. Sejnowski. Associative memory and hippocampal place cells. International Journal of Neural Systems, Supplementary Issue, 1995, page 81–86, 1995.

28. J. O’Keefe and N. Burgess. Dual phase and rate coding in hippocampal place cells: Theoretical significance and relationship to entorhinal grid cells. Hippocampus, 15(7):853–866, 2005. doi: 10.1002/hipo.20115.

29. T. Solstad, E. I. Moser, and G. T. Einevoll. From grid cells to place cells: A mathematical model. Hippocampus, 16(12): 1026–1031, 2006. doi: 10.1002/hipo.20244.

30. M. Fyhn, T. Hafting, A. Treves, M. B. Moser, and E. I. Moser. Hippocampal remapping and grid realignment in entorhinal cortex. Nature, 446(7132):190–194, 2007. doi: 10.1038/nature05601.

31. J. D. Monaco and L. F. Abbott. Modular realignment of entorhinal grid cell activity as a basis for hippocampal remapping. Journal of Neuroscience, 31(25):9414–9425, 2011. doi: 10.1523/Jneurosci.1433-11.2011.

32. T. Hainmueller, A. Cazala, L. W. Huang, and M. Bartos. Subfield-specific interneuron circuits govern the hippocampal response to novelty in male mice. Nature Communications, 15(1), 2024. doi: ARTN714 10.1038/s41467-024-44882-3.

33. B. R. Kanter, C. M. Lykken, D. Avesar, A. Weible, J. Dickinson, B. Dunn, N. Z. Borgesius, Y. Roudi, and C. G. Kentros. A novel mechanism for the grid-to-place cell transformation revealed by transgenic depolarization of medial entorhinal cortex layer ii. Neuron, 93(6):1480–1492 e6, 2017. ISSN 1097-4199 (Electronic) 0896-6273 (Linking). doi: 10.1016/j.neuron.2017.03.001. Kanter, Benjamin R Lykken, Christine M Avesar, Daniel Weible, Aldis Dickinson, Jasmine Dunn, Benjamin Borgesius, Nils Z Roudi, Yasser Kentros, Clifford G eng R01 MH097130/MH/NIMH NIH HHS/2017/03/24 Neuron. 2017 Mar 22;93(6):1480–1492.e6. doi: 10.1016/j.neuron.2017.03.001.

34. C. M. Lykken, B. R. Kanter, A. Nagelhus, J. Carpenter, M. Guardamagna, E. I. Moser, and M.-B. Moser. Functional independence of entorhinal grid cell modules enables remapping in hippocampal place cells. bioRxiv, 2025. doi: 10.1101/2025.09.24.677985.

35. A. Samsonovich and B. L. McNaughton. Path integration and cognitive mapping in a continuous attractor neural network model. Journal of Neuroscience, 17(15):5900–5920, 1997.

36. C. Dong, A. D. Madar, and M. E. J. Sheffield. Distinct place cell dynamics in ca1 and ca3 encode experience in new environments. Nature Communications, 12(1), 2021. doi: ARTN2977 10.1038/s41467-021-23260-3.

37. David Dupret, Joseph O’Neill, Barty Pleydell-Bouverie, and Jozsef Csicsvari. The reorganization and reactivation of hippocampal maps predict spatial memory performance. Nature Neuroscience, 13(8):995–1002, 2010. doi: 10.1038/nn.2599.

38. Isha Zutshi, Manuel Valero, Antonio Fernández-Ruiz, and György Buzsáki. Extrinsic control and intrinsic computation in the hippocampal ca1 circuit. Neuron, 110(4):658–673.e5, 2022. doi: 10.1016/j.neuron.2021.11.015.

39. K. C. Bittner, C. Grienberger, S. P. Vaidya, A. D. Milstein, J. J. Macklin, J. Suh, S. Tonegawa, and J. C. Magee. Conjunctive input processing drives feature selectivity in hippocampal ca1 neurons. Nature Neuroscience, 18(8):1133–+, 2015. doi: 10.1038/nn.4062.

40. K. C. Bittner, A. D. Milstein, C. Grienberger, S. Romani, and J. C. Magee. Behavioral time scale synaptic plasticity underlies ca1 place fields. Science, 357(6355):1033–1036, 2017. doi: 10.1126/science.aan3846.

41. S. Blankvoort, M. P. Witter, J. Noonan, J. Cotney, and C. Kentros. Marked diversity of unique cortical enhancers enables neuron-specific tools by enhancer-driven gene expression. Current Biology, 28(13):2103–+, 2018. doi: 10.1016/j.cub.2018.05.015.

42. A. S. Chuong, M. L. Miri, V. Busskamp, G. A. Matthews, L. C. Acker, A. T. Sorensen, A. Young, N. C. Klapoetke, M. A. Henninger, S. B. Kodandaramaiah, M. Ogawa, S. B. Ramanlal, R. C. Bandler, B. D. Allen, C. R. Forest, B. Y. Chow, X. Han, Y. Lin, K. M. Tye, B. Roska, J. A. Cardin, and E. S. Boyden. Noninvasive optical inhibition with a red-shifted microbial rhodopsin. Nat Neurosci, 17(8):1123–9, 2014. ISSN 1546-1726 (Electronic) 1097-6256 (Linking). doi: 10.1038/nn.3752.

43. M. Guardamagna, O. Chadney, F. Stella, Q. W. Zhang, C. Kentros, and F. P. Battaglia. Direct entorhinal control of ca1 temporal coding. Nature Communications, 16(1), 2025. doi: ARTN6430 10.1038/s41467-025-61453-2.

44. K. D. Harris, H. Hirase, X. Leinekugel, D. A. Henze, and G. Buzsáki. Temporal interaction between single spikes and complex spike bursts in hippocampal pyramidal cells. Neuron, 32(1):141–149, 2001. doi: Doi10.1016/S0896-6273(01)00447-0.

45. N. Schmitzer-Torbert, J. Jackson, D. Henze, K. Harris, and A. D. Redish. Quantitative measures of cluster quality for use in extracellular recordings. Neuroscience, 131(1):1–11, 2005. doi: 10.1016/j.neuroscience.2004.09.066.

46. Stefan Leutgeb, Jill K. Leutgeb, Carol A. Barnes, Edvard I. Moser, Bruce L. McNaughton, and May-Britt Moser. Independent codes for spatial and episodic memory in hippocampal neuronal ensembles. Science, 309(5734):619–623, 2005. doi: 10.1126/science.1114037.

47. Christine Grienberger, Xiaowei Chen, and Arthur Konnerth. Nmda receptor-dependent multidendrite ca2+ spikes required for hippocampal burst firing in vivo. Neuron, 81(6):1274–1281, 2014. doi: 10.1016/j.neuron.2014.01.014.

48. J. B. Priestley, J. C. Bowler, S. Rolotti, S. Fusi, and A. Losonczy. Article signatures of rapid plasticity in hippocampal ca1 representations during novel experiences. Neuron, 110(12):1978–+, 2022. doi: 10.1016/j.neuron.2022.03.026.

49. Eunji Kong, Erfan Zabeh, Zhenrui Liao, Tiberiu S. Mihaila, Caroline Wilson, Charan Santhirasegaran, Darcy S. Peterka, Attila Losonczy, and Tristan Geiller. Recurrent connectivity shapes spatial coding in hippocampal ca3 subregions. bioRxiv, page 2024.11.07.622379, 2024. doi: 10.1101/2024.11.07.622379.

50. H. Dvorak-Carbone and E. M. Schuman. Long-term depression of temporoammonic-ca1 hippocampal synaptic transmission. Journal of Neurophysiology, 81(3):1036–1044, 1999. doi: DOI 10.1152/jn.1999.81.3.1036.

51. M. Remondes and E. M. Schuman. Direct cortical input modulates plasticity and spiking in ca1 pyramidal neurons. Nature, 416(6882):736–740, 2002. doi: DOI 10.1038/416736a.

52. M. E. J. Sheffield, M. D. Adoff, and D. A. Dombeck. Increased prevalence of calcium transients across the dendritic arbor during place field formation. Neuron, 96(2):490–+, 2017. doi: 10.1016/j.neuron.2017.09.029.

53. L. Z. Fan, D. K. Kim, J. H. Jennings, H. Tian, P. Y. Wang, C. Ramakrishnan, S. Randles, Y. J. Sun, E. Thadhani, Y. S. Kim, S. Quirin, L. Giocomo, A. E. Cohen, and K. Deisseroth. All-optical physiology resolves a synaptic basis for behavioral timescale plasticity. Cell, 186(3):543–+, 2023. doi: 10.1016/j.cell.2022.12.035.

54. K. C. Gonzalez, A. Negrean, Z. R. Liao, S. Terada, G. F. Zhang, S. M. Lee, K. Ocsai, B. J. Rozsa, M. Z. Lin, F. Polleux, and Losonczy. Synaptic basis of feature selectivity in hippocampal neurons. Nature, 637(8048), 2025. doi: 10.1038/s41586-024-08325-9.

55. J. K. O’Hare, J. M. Wang, M. D. Shala, F. Polleux, and A. Losonczy. Distal tuft dendrites predict properties of new hippocampal place fields. Neuron, 113(12):1969–1982, 2025. doi: 10.1016/j.neuron.2025.03.029.

56. M. Megías, Z. Emri, T. F. Freund, and A. I. Gulyás. Total number and distribution of inhibitory and excitatory synapses on hippocampal ca1 pyramidal cells. Neuroscience, 102(3):527–540, 2001. doi: Doi10.1016/S0306-4522(00)00496-6.

57. T. Takeuchi, A. J. Duszkiewicz, A. Sonneborn, P. A. Spooner, M. Yamasaki, M. Watanabe, C. C. Smith, G. Fernández, K. Deisseroth, R. W. Greene, and R. G. M. Morris. Locus coeruleus and dopaminergic consolidation of everyday memory. Nature, 537(7620):357–+, 2016. doi: 10.1038/nature19325.

58. A. M. Kaufman, T. Geiller, and A. Losonczy. A role for the locus coeruleus in hippocampal ca1 place cell reorganization during spatial reward learning. Neuron, 105(6):1018–+, 2020. doi: 10.1016/j.neuron.2019.12.029.

59. M. Udakis, V. Pedrosa, S. E. L. Chamberlain, C. Clopath, and J. R. Mellor. Interneuron-specific plasticity at parvalbumin and somatostatin inhibitory synapses onto ca1 pyramidal neurons shapes hippocampal output. Nature Communications, 11(1), 2020. doi: ARTN4395 10.1038/s41467-020-18074-8.

60. J. Palacios-Filardo, M. Udakis, G. A. Brown, B. G. Tehan, M. S. Congreve, P. J. Nathan, A. J. H. Brown, and J. R. Mellor. Acetylcholine prioritises direct synaptic inputs from entorhinal cortex to ca1 by differential modulation of feedforward inhibitory circuits (sep, 10.1038/s41467-021-25280-5, 2021). Nature Communications, 12(1), 2021. doi: ARTN7265 10.1038/s41467-021-27351-z.

61. M. Udakis, M. D. B. Claydon, H. W. Zhu, E. C. Oakes, and J. R. Mellor. Hippocampal olm interneurons regulate ca1 place cell plasticity and remapping. Nature Communications, 16(1), 2025. doi: ARTN9912 10.1038/s41467-025-64859-0.

62. B. R. Rost, J. Wietek, O. Yizhar, and D. Schmitz. Optogenetics at the presynapse. Nature Neuroscience, 25(8):984–998, 2022. doi: 10.1038/s41593-022-01113-6.

63. M. I. Schlesiger, B. L. Boublil, J. B. Hales, J. K. Leutgeb, and S. Leutgeb. Hippocampal global remapping can occur without input from the medial entorhinal cortex. Cell Reports, 22(12):3152–3159, 2018. doi: 10.1016/j.celrep.2018.02.082.

64. J. C. Bowler and A. Losonczy. Direct cortical inputs to hippocampal area ca1 transmit complementary signals for goal-directed navigation. Neuron, 111(24):4071–4085 e6, 2023. ISSN 1097-4199 (Electronic) 0896-6273 (Print) 0896-6273 (Linking). doi: 10.1016/j.neuron.2023.09.013.

65. M. Guardamagna, R. Eichler, R. Pedrosa, A. Aarts, A. F. Meyer, and F. P. Battaglia. The hybrid drive: a chronic implant device combining tetrode arrays with silicon probes for layer-resolved ensemble electrophysiology in freely moving mice. Journal of Neural Engineering, 19(3), 2022. doi: ARTN036030 10.1088/1741-2552/ac6771.

66. J. Voigts, J. P. Newman, M. A. Wilson, and M. T. Harnett. An easy-to-assemble, robust, and lightweight drive implant for chronic tetrode recordings in freely moving animals. Journal of Neural Engineering, 17(2), 2020. doi: ARTN026044 10.1088/1741-2552/ab77f9.

67. J. H. Siegle, A. C. López, Y. A. Patel, K. Abramov, S. Ohayon, and J. Voigts. Open ephys: an open-source, plugin-based platform for multichannel electrophysiology. Journal of Neural Engineering, 14(4), 2017. doi: ARTN045003 10.1088/1741-2552/aa5eea.

68. William E. Skaggs, Bruce L. McNaughton, Matthew A. Wilson, and Carol A. Barnes. Theta phase precession in hippocampal neuronal populations and the compression of temporal sequences. Hippocampus, 6(2):149–172, 1996. doi: 10.1002/(SICI)1098-1063(1996)6:2<149::AID-HIPO6>3.0.CO;2-K.

